# *Tbx1*, a 22q11.2-encoded gene, is a link between alterations in fimbria myelination and cognitive speed in mice

**DOI:** 10.1101/2021.03.29.437581

**Authors:** Takeshi Hiramoto, Akira Sumiyoshi, Takahira Yamauchi, Kenji Tanigaki, Qian Shi, Gina Kang, Rie Ryoke, Hiroi Nonaka, Shingo Enomoto, Takeshi Izumi, Manzoor A. Bhat, Ryuta Kawashima, Noboru Hiroi

**Author notes:** Corresponding author Noboru Hiroi, PhD, Department of Pharmacology; Department of Cellular and Integrative Physiology; Department of Cell Systems and Anatomy, University of Texas Health Science Center, San Antonio Room 211B, 7703 Floyd Curl Drive San Antonio, TX 78229 1. These authors contributed equally to the work (Takeshi Hiramoto, Akira Sumiyoshi, Takahira Yamauchi).

## Abstract

Copy number variants (CNVs) have provided a reliable entry point to identify structural correlates of atypical cognitive development. Hemizygous deletion of human chromosome 22q11.2 is associated with impaired cognitive function; however, the mechanisms by which numerous genes encoded in this CNV contribute to cognitive deficits via diverse structural alterations in the brain remain unclear. This study aimed to determine the cellular basis of the link between alterations in brain structure and cognitive functions in a mouse model. The heterozygosity of *Tbx1, a* 22q11.2 gene, altered the composition of myelinated axons in the fimbria, reduced oligodendrocyte production capacity, and slowed the acquisition of spatial memory and cognitive flexibility. Our findings provide a cellular basis for specific cognitive dysfunctions that occur in patients with loss-of-function *TBX1* variants and 22q11.2 hemizygous deletion.

**Teaser:** A risk gene for autism alters myelin composition in the hippocampal connection and slows cognitive speed.

## INTRODUCTION

Although copy number variants (CNVs) are rare and occur in <1% of patients with any psychiatric disorder, they are robustly and consistently associated with developmental neuropsychiatric disorders (*1, 2*). Moreover, CNVs affect specific cognitive functions independent of clinically defined mental illness (*3*). Currently available pharmaceutical medications do not significantly improve cognitive deficits associated with many mental disorders due to a lack of understanding of their causative mechanistic targets.

Despite their robust association with cognitive impairments and psychiatric disorders, CNVs pose a challenge when attempting to understand the composition of contributory genes, as accurate identification of CNV-encoded genes contributing to human phenotypes remains difficult. A recent large-scale genome-wide exome screening study reported that protein-truncating variants of genes encoded in several large-sized CNVs are linked with autism spectrum disorder (ASD) (*4*). However, failure to detect similar variants of other CNV-encoded genes may be attributable to their rarity, as larger sample sizes enable the identification of more gene variants than smaller-scale analyses(*4, 5*). Moreover, variants in promoters and enhancers may contribute to phenotypes (*6*). Variants of CNV-encoded single genes may simply not exist, and single-gene hemizygosity or duplication, as part of a CNV, may play the role of a driver gene. Thus, there is a need to utilize complementary approaches to identify driver genes encoded by large CNVs.

There are more human and mouse studies of human chromosome 22q11.2 deletion than other CNVs, given that it was found to be associated with mental illness much earlier than other CNVs (*7*). Hemizygous deletion of 22q11.2 CNVs is robustly associated with diverse disorders, including ASD, attention-deficit/hyperactivity disorder, schizophrenia, and intellectual disability (ID) (*8*). Moreover, individuals with 22q11.2 hemizygosity exhibit deterioration in specific cognitive functions, including the accuracy and speed of memory acquisition, executive functions, and social cognition (*9–11*). Further, cognitive impairment precedes and predicts the onset of schizophrenia among 22q11.2 hemizygosity carriers (*12, 13*). In addition, recent large-scale imaging studies have demonstrated altered white matter integrity in the brains of 22q11.2 hemizygous deletion carriers (*14–16*); no DTI-MRI analysis of mouse models of 22q11.2 hemizygosity have been reported. However, since many regions show altered white matter integrity and this CNV contains a minimum of 30 protein-coding genes, the exact causative associations among encoded genes, structural alterations, and atypical cognitive development remain unclear.

Rare loss-of-function variants (e.g., frameshift deletion) of *TBX1,* a gene encoded by a 22q11.2 CNV, have been associated with ASD, ID, and seizures (*17–20*). However, these *TBX1* variant carriers also exhibit single nucleotide variants (SNVs) in other genes (*19*), and only a few cases/families with those variants have been identified. The causative structural substrates in the brain mediating the impacts of *Tbx1* deficiency on cognitive impairment remain unknown.

Mouse studies have provided a complementary means to address limitations of these human studies by systematically examining the roles of small chromosomal segments and individual genes in behaviors against a homogeneous genetic background (*8, 20–30*). These studies have demonstrated that some, but not all, 22q11.2-encoded single genes contribute to select behavioral targets (*8, 26, 29*). For example, our results have revealed that *Tbx1* heterozygosity impairs social interaction and communication (*24, 27, 28, 30*). The present study aimed to determine the structural and cellular basis underlying the effects of *Tbx1* heterozygosity on specific cognitive functions in a congenic mouse model.

## RESULTS

There are alterations in white matter microstructures in many brain regions of 22q11.2 hemizygosity carriers (*14–16*). However, little is known regarding the exact nature of altered white matter microstructures and driver genes that affect both the white matter and cognitive functions.

### Analysis of white matter structures

#### *Tbx1* deficiency decreases fractional anisotropy (FA) signals in the fimbria

*Tbx1* +/- mice and their +/+ littermates underwent *ex vivo* diffusion tensor imaging (DTI)- magnetic resonance imaging (MRI). We analyzed 19 brain regions (Fig. S1), as defined by the standard regional classification of the mouse brain (*31*). The FA value is the most histologically validated DTI-MRI metric (*32*). However, since FA signals of <0.3 are not reliably correlated with the degree of myelination (*33*) and cannot be accurately aligned across individual animals (*34*), we selected regions with FA values ≥0.3. The corpus callosum, anterior commissure, internal capsule, and fimbria met this criterion (**Fig. S2A**). The fimbria was the only region exhibiting a significant change: FA values were lower in +/- mice than in +/+ mice (**Fig. 1**). Consistently, the fimbria exhibited the largest effect size for genotype-dependent differences in FA values (**Fig. S2B**). There were no significant differences in axial diffusivity (AD), radial diffusivity (RD), or mean diffusivity (MD) values between +/+ and +/- mice (**Fig. S3-5**).

**Figure 1.**
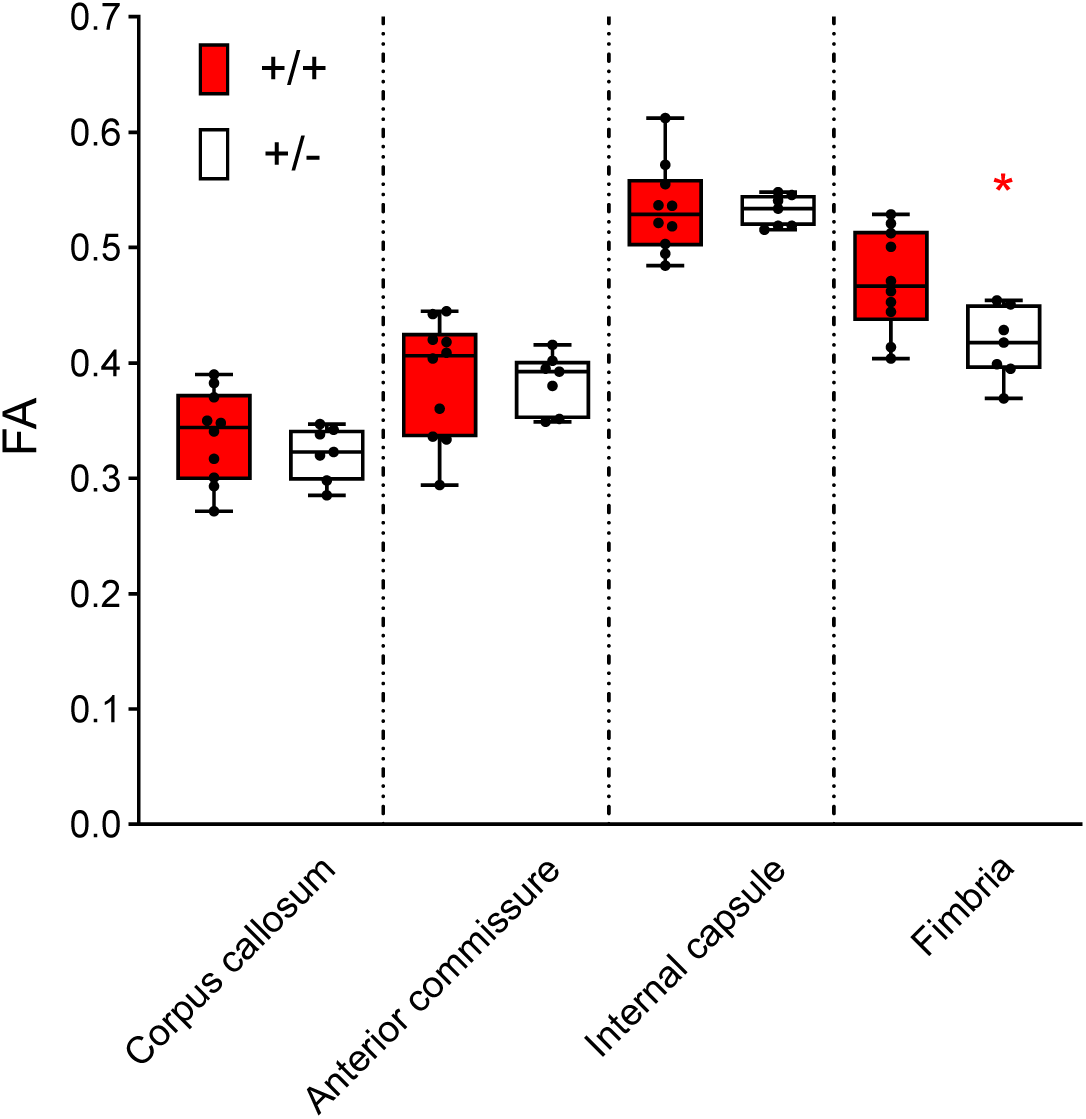
Box-and-Whisker plots of fractional anisotropy (FA) values of the four regions with FA >0.3. Analysis using a generalized linear mixed model revealed a region- dependent differential effect of genotype on FA values (Genotype x Region, F(3,45) = 7.337, P < 0.001). Mann–Whitney U-tests revealed a significant between-genotype difference in the fimbria only (*, U = 11, p = 0.0185). +/+, n = 10; +/-, n = 7.

#### *Tbx1* deficiency reduces myelination in the fimbria

DTI-MRI analysis of the mouse brain has limited spatial resolution, as well as technical and interpretative limitations (*32, 35*). The structural classifications of Ma et al. (*31*) include the fimbria, fornix, stria terminalis, and hippocampal commissure in the “fimbria”. To circumvent these limitations and histologically validate the DTI-MRI findings, we used the non-hydroscopic gold-phosphate complex Black-Gold II (*36*). This method provides more consistent staining than hydroscopic gold chloride staining and higher contrast and resolution than lipid soluble dyes (e.g., Luxol Fast Blue). Black-Gold II also directly stains myelin, unlike markers of myelin components (e.g., myelin basic protein [MBP]), which may not perfectly correlate with the degree of myelination. We examined regions with the largest and second largest effect sizes among FA values ≥0.3: the fimbria and corpus callosum (**Fig. S2B; S6**). The intensity of gold staining was lower in the anterior fimbria of +/- mice than in that of +/+ mice (**Fig. 2A**). There was no statistically detectable between-genotype difference in the posterior fimbria or anterior/posterior corpus callosum (**Fig. 2B-D**).

**Figure 2.**
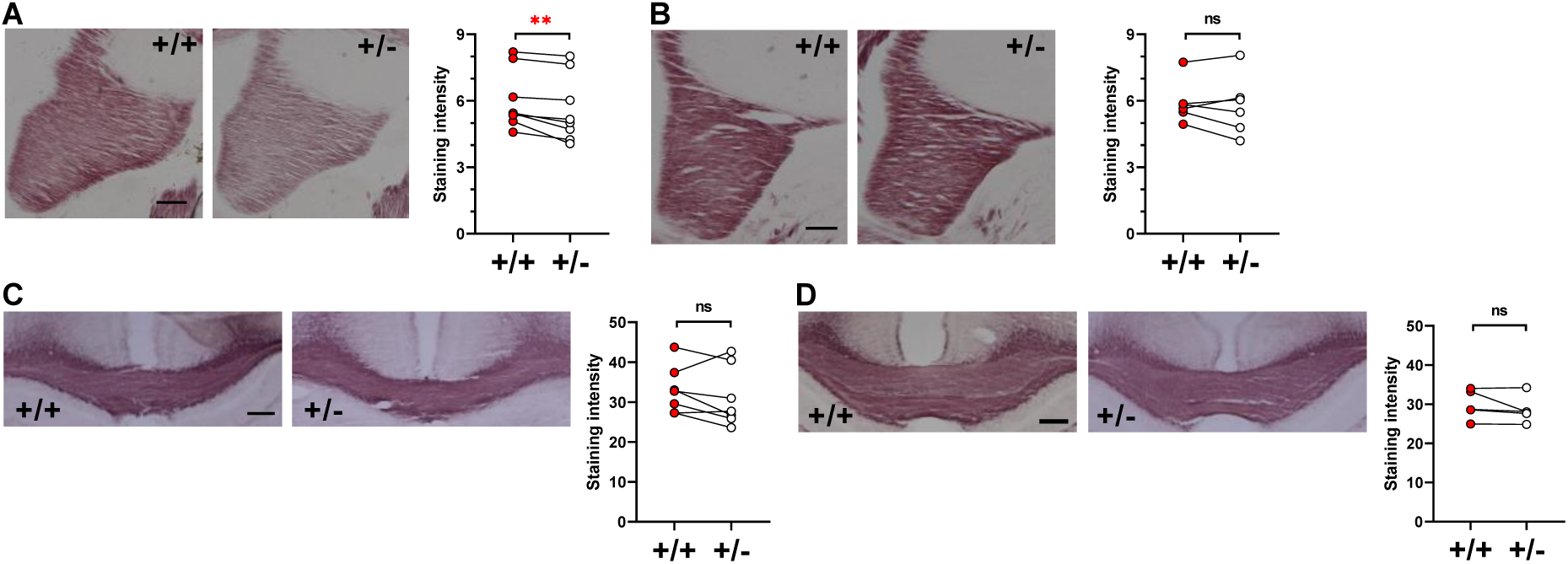
Black-Gold II staining of myelin in the anterior fimbria (**A**), posterior fimbria (**B**), anterior corpus callosum (**C**), and posterior corpus callosum (**D**) (see **Supplemental Fig. S6**). Representative images of gold-stained myelin (left panels) and staining intensities of each pair of +/+ and +/- mice (right panels) are shown. Given that the assumptions of normality and homogeneity of variance of the raw data from all sections were not met, we used non-parametric Wilcoxon tests for paired +/+ and +/- sections of comparable anterior-posterior positions within each slide. Compared with +/+ littermates, +/- mice exhibited significantly decreased levels of gold staining in the anterior fimbria (**, p = 0.0078), but not in the posterior fimbria (not significant (ns), p = 0.5625), anterior corpus callosum (ns, p = 0.2969), or posterior corpus callosum (ns, p = 0.1875). Anterior fimbria, 8 +/+ mice and 8 +/- mice; posterior fimbria, 6 +/+ mice and 6 +/- mice; anterior corpus callosum, 7 +/+ mice and 7 +/- mice; posterior corpus callosum, 5 +/+ mice and 5 +/- mice. Scale bar = 200 μm.

#### *Tbx1* deficiency reduces large myelinated axons in the fimbria

We used electron microscopy (EM) to characterize the myelination of axons in the fimbria and corpus callosum at the ultrastructural level. Myelination appeared thicker and thinner in the fimbria and corpus callosum, respectively, of +/- mice than in that of +/+ mice (**Fig. 3AB**). We compared g-ratios (i.e., the ratio of axon diameter to the axon + myelin diameter) to quantitatively evaluate relative myelin thickness (**Fig. 3C, D**). The g-ratios of +/+ mice plateaued slightly above 0.8, which is an expected value for the optimal efficiency of axon myelination in the central nervous system (CNS) (*37*). The g-ratios of +/- mice were smaller (i.e., relatively thicker myelin sheath) in the fimbria (**Fig. 3C**), but not in the corpus callosum (**Fig. 3D**). Volume is a limiting factor in the CNS: The myelination efficiency steeply decreases when myelin thickness deviates from the optimal g-ratio (∼0.8) (*37*) regardless of whether it is hyper- or hypo-myelination. Therefore, this gene deficiency results in a functionally sub-optimal population of axons in the fimbria.

**Figure 3.**
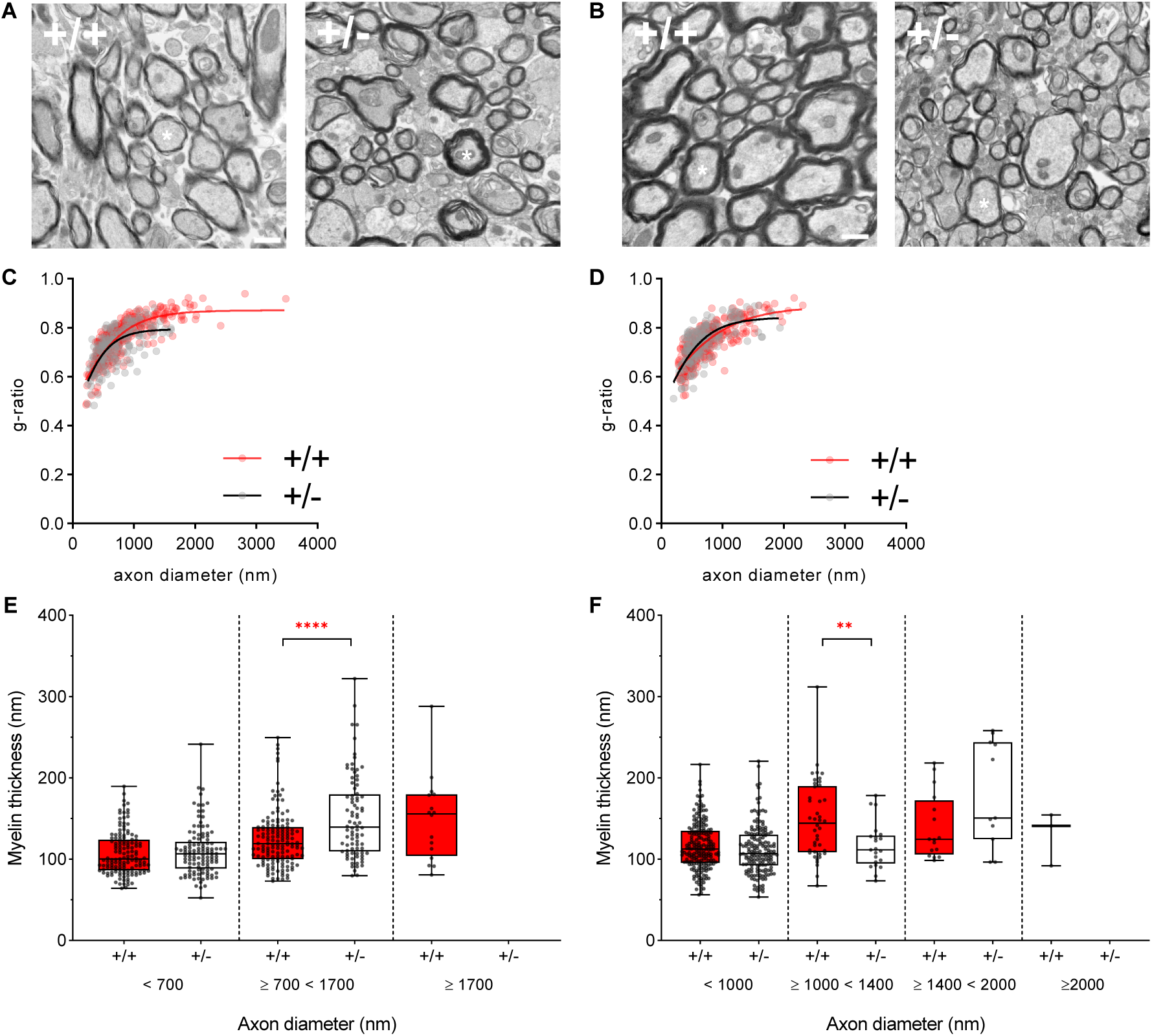
Electron microscopy (EM) images of myelin in the fimbria (**A**) and corpus callosum (**B**). We analyzed 300 and 200 axons in the fimbria of both hemispheres in three +/+ and two +/- mice, respectively. We analyzed 260 and 200 axons in the corpus callosum of both hemispheres in +/+ and +/- mice, respectively. Ten images were obtained from the fimbria or corpus callosum of each mouse, except for one +/+ mouse that had six available images of the corpus callosum. Ten randomly chosen myelinated axons were analyzed from each image. Data from each image and axon were treated as random duplicates. Scale bar = 800nm. (**C**) G-ratios in the fimbria across captured images and axons within each image were consistently lower in +/- mice than in +/+ mice. Since the normality assumptions were not met (+/+, p = 0.003; +/-, p = 0.005), we used a generalized linear mixed model (Genotype, F(1,300) = 19.539, p < 0.001; Genotype x Image, F(9,300) = 1.011, p = 0.431; Genotype x Axon, F(9,300) = 0.519, p = 0.861; Genotype x Image x Axon, F(81,300) = 1.211, p = 0.129). **D**) G-ratios in the corpus callosum did not differ between the genotypes (Genotype, F(1,260) = 0.025, p=0.876; Genotype x Axon, F(9,260) = 0.760, p = 0.654; Genotype x Image x Axon, F(81,260) = 1.009, p = 0.467). The g-ratio was calculated as g = d/D, where d and D represent the axon and axon + myelin diameters, respectively. (**E,F**) Box-and-Whisker plots of myelin thickness (Y, nm) for a range of axon diameters (X, nm) in the fimbria (**E**) and corpus callosum (**F**). The ranges are based on segments where +/+ and +/- differed (see **Fig. S7AB**). In +/- mice, myelin thickness in the fimbria was increased for axon diameters ≥700 nm and <1,700 nm (Mann–Whitney U = 4,202, p < 0.0001,***) (see white stars in **A**), but not for those with diameters <700 nm (Mann–Whitney U = 7,653, p = 0.5473). There were no fimbria axons with diameters ≥1700 nm in +/- mice (see **Fig. S8; Table S1**). In +/- mice, myelin thickness in the corpus callosum was decreased for axons with diameters ≥1,000 nm and <1,400 nm (Mann–Whitney U = 258, p = 0.0071, **) but not for those with other diameter ranges (<1,000 nm; Mann–Whitney U =14,989, p = 0.0881; ≥1,400 nm; Mann-Whitney U =56, p = 0.1214). There were no corpus callosum axons with diameters >1,400 nm axons in +/- mice.

A further in-depth analysis revealed that 1) myelin was thicker in axons with diameters ≥700 nm to <1,700nm in the fimbria of +/- mice than of +/+ mice (**Fig. 3E**; **Fig. S7A**); 2) there were proportionally fewer myelinated axons ≥1,200 nm in diameter in the fimbria of +/- mice than of +/+ mice (**Fig. S8A**; **S-Table 1**); and 3) there were no myelinated axons ≥1,700 nm in diameter in the fimbria of +/- mice (**Fig. 3E; S8A**). In the corpus callosum, myelin was thinner in axons with diameters between 1,000 nm and 1,400 nm in +/- mice than in +/+ mice (**Fig. 3F**). The relative proportion of axons in the corpus callosum was increased in axons between ≥400 nm and <800 nm in diameter and was decreased in axons ≥800 nm in diameter in +/- mice (**Fig. S8B**; **S-Table 1**). There were no myelinated corpus callosum axons with diameters ≥2,000 nm in +/- mice (**Fig. 3F**; **Fig. S8B**).

In sum, *Tbx1* heterozygous mice lacked large (≥1,700 nm) myelinated axons and exhibited hyper-myelination of axons up to 1,700 nm in diameter in the fimbria. In the corpus callosum of +/- mice, axons with diameters ranging from 1,000 to 1,400 nm were hypomyelinated. In addition, +/- mice exhibited proportionally more myelinated axons with diameters between 400 nm and 800 nm but less myelinated axons with diameters ≥800 nm in the corpus callosum.

#### *Tbx1* heterozygosity impacts a molecule critical for early oligodendrogenesis

Oligodendrocytes and their precursor cells are present locally in the fimbria and, to a lesser extent, in the corpus callosum. The molecular steps through which *Tbx1* impacts oligodendrogenesis and myelination remains unknown. Given that *Tbx1* mRNA is reduced in the fimbria and corpus callosum of *Tbx1* +/- mice compared to +/+ mice (**Fig. 4**), we examined the impact of a dose reduction of *Tbx1* mRNA on molecules functionally critical for each step of oligodendrogenesis and myelination in 2- to 3-month-old *Tbx1* +/- and +/+ littermates, using qRT-PCR. The myelinating process of the fimbria starts in the second neonatal week and reaches its peak around postnatal day 24-37 in rodents (*38*).

**Figure 4.**
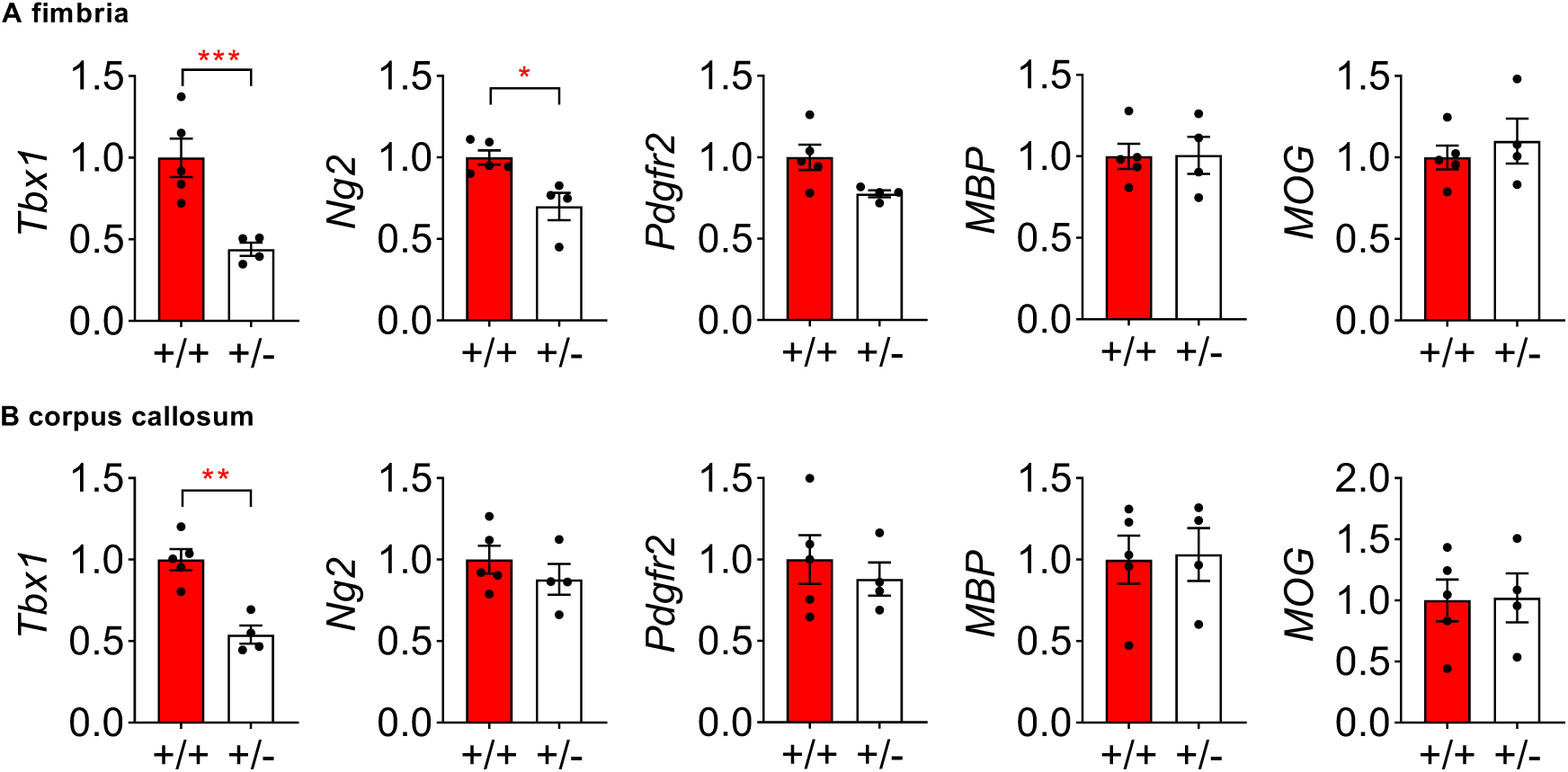
Relative mRNA expression levels (mean±standard error of the mean [SEM]) for *Tbx1*, *Ng2*, *Pdgfr2*, myelin basic protein (MBP), and myelin oligodendrocyte glycoprotein (MOG) in the fimbria (**A**) and corpus callosum (**B**) of *Tbx1*+/+ (n = 5) and +/- (n = 4) mice. *Tbx1* mRNA levels were lower in the fimbria (**A**, t(7)=4.081, p = 0.0047, ***) and corpus callosum (**B**, t(7)=5.221, p = 0.0012, **) of +/- mice than in those of +/+ mice. In the fimbria, levels of *Ng2* (**A**, t(7)=3.394, p = 0.0115, *) were lower in +/- mice than in +/+ mice. These significant differences survived Benjamini–Hochberg’s correction at the false discovery rate (FDR) of 5%. There were no other significant differences in the fimbria or corpus callosum (**A**,**B**, p>0.05).

Ng2 (Cspg4) and Pdfgr2, markers of oligodendrocyte precursor cells, are functionally required for the production of oligodendrocyte precursor cells (*39, 40*). We found that mRNA levels of *Ng2* and, but not of *Pdgrf2,* were selectively lower in the fimbria of +/- mice than in that of +/+ mice (**Fig. 4A**). There was no detectable difference in *Ng2* or *Pdgfr2* mRNA levels in the corpus callosum between +/+ and +/- mice (**Fig. 4B**). MBP is essential for the maintenance of myelin and is involved in the adhesion and compaction of the cytosolic membrane leaflets that form the structural basis of multilayered myelin (*41, 42*). Myelin oligodendrocyte glycoprotein (MOG) is a marker of mature oligodendrocytes and myelin, although it is not functionally critical for myelin formation or maintenance (*43*). No differences in MBP or MOG levels were observed in the fimbria or corpus callosum between +/+ and +/- mice (**Fig. 4AB**). This *in vivo* analysis indicated that *Tbx1* heterozygosity selectively impacts the very early molecular step of oligodendrogenesis locally in the fimbria but has no effect on molecules required for myelin formation and maintenance in this location.

#### *Tbx1* heterozygosity reduces oligodendrocyte generation

Another source of oligodendrocytes in the fimbria is the population of adult neural progenitor cells in the subventricular zone (*44–47*), which is distinct from those generating neurons (*48*). Given the enrichment of Tbx1 protein in the adult subventricular zone (SVZ) (*24*), we aimed to determine whether *Tbx1* heterozygosity affects the cell- autonomous capacity of this oligodendrocyte population. Progenitor cells were taken from the lateral ventricular wall, including the subventricular zone, of 3-week-old *Tbx1* +/- and +/+ littermates and cultured and differentiated into oligodendrocytes *in vitro*. Progenitor cells derived from the subventricular zone of *Tbx1* +/- mice produced fewer O4-positive immature and mature oligodendrocytes than those derived from +/+ mice (**Fig. 5AB**). This *in vitro* assay demonstrated that *Tbx1* heterozygosity reduced oligodendrocyte production from progenitor cells of the SVZ in a cell-autonomous manner.

**Figure 5.**
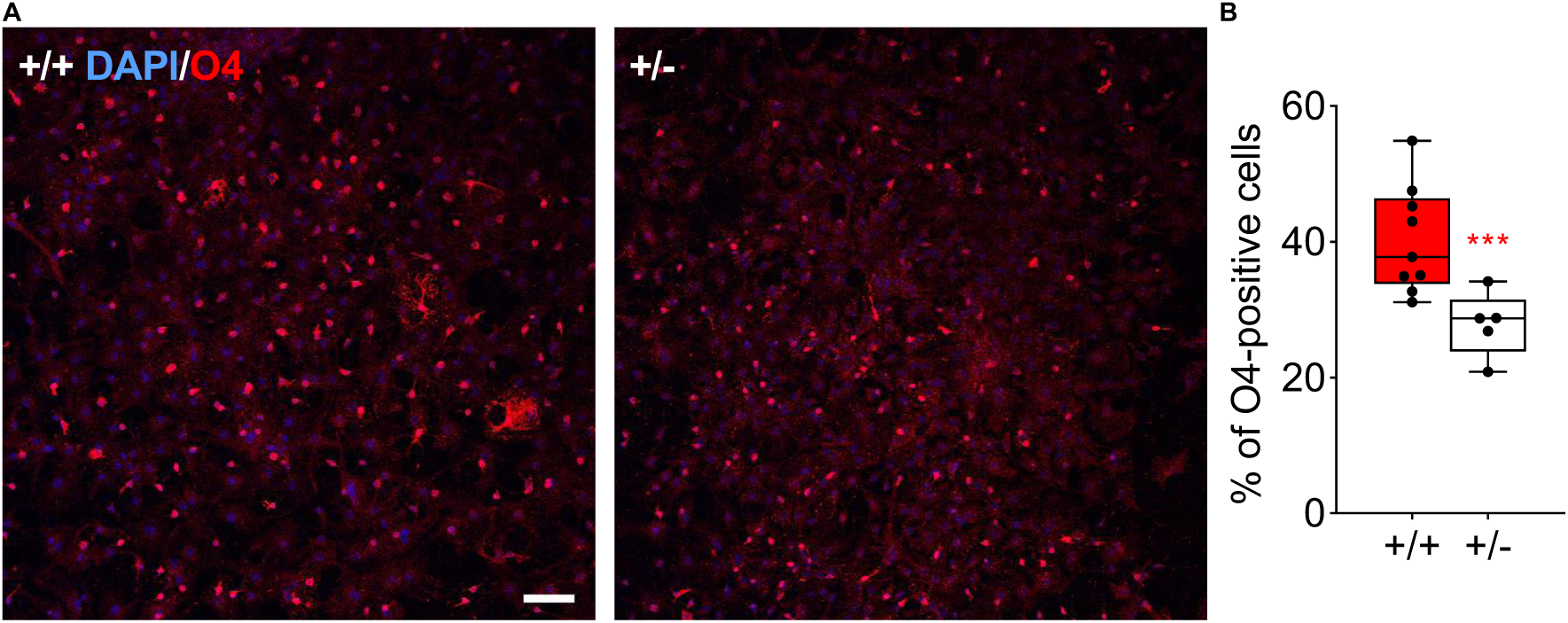
Representative images (**A**) and Box-and-Whisker plots (**B**) of O4-positive (red) oligodendrocytes among all DAPI-positive (blue) cells in culture. Since the assumption of normality was not met (Shapiro–Wilk tests: +/+, W(35) = 0.888, p = 0.002; +/-, W(19) = 0.898, p = 0.045), we applied a generalized linear mixed model of log transformed data. Progenitor cells derived from the lateral ventricular walls of P21 *Tbx1* +/- mice produced consistently fewer O4-positive oligodendrocytes than those of +/+ mice across the cultures (Genotype, F(1,11.451) = 12.841. p = 0.004, ***; Image field, F(3, 33.978) = 0.609, p = 0.614; Genotype x Image field, F(3,33.978) = 0.134, p = 0.939). Scale bar = 200 μm. +/+, n = 9; +/-, n = 5.

DTI-MRI analysis detected the region exhibiting the most robust alteration in white matter integrity. This observation was validated using Black-Gold II staining, which provided a higher anatomical resolution of the net myelination levels. Although decreased FA values and reduced net myelin signals are suggestive of less myelin, axonal degeneration, reduced axonal density, or changes in axonal organization (*49*), our EM analyses complemented these assessments of the net signal intensities by demonstrating a loss of large myelinated axons in the fimbria and reduced myelination in axons with specific diameters in the corpus callosum. Our qRT-PCR and *in vitro* analyses further indicated that *Tbx1* heterozygosity reduced levels of the molecule needed for the generation of local oligodendrocyte precursor cells and the production capacity of oligodendrocytes in the lateral ventricular wall.

### Analysis of cognitive functions

Individuals with 22q11.2 hemizygous deletions exhibit lower scores on measures of attention, executive function, processing speed, visual memory, visuospatial skills, and social cognition (*9, 11*). However, the link between structural alterations caused by single 22q11.2 genes and changes in cognitive function remains unknown. In addition, although human studies have reported an association between loss-of-function *TBX1* variants and developmental neuropsychiatric disorders (*17–20*), their effects on cognitive function remain uncharacterized. Since we observed that *Tbx1* heterozygosity leads to myelin alterations in the fimbria, we examined its effects on cognitive capacities known to rely on the fimbria.

#### *Tbx1* heterozygosity slows the acquisition of spatial reference memory

The spatial reference memory version of the Morris water maze requires an intact fimbria, whereas the visual cued version depends on the dorsal striatum in rodents (*50, 51*). Humans with 22q11.2 hemizygosity exhibit impaired spatial processing and memory (*9, 52–54*). Although a previous study reported that spatial reference memory retention and recall in the Morris water maze were normal in a mouse model of 22q11.2 hemizygosity (*55*), no studies have investigated these capacities in the acquisition phase.

*Tbx1* +/- mice exhibited delayed spatial memory acquisition in the Morris water maze (**Fig. 6A-C**). In contrast, there was no between-genotype difference in the re-probe test (**Fig. 6D**) or during visual cue memory acquisition (**Fig. 6E**). These data indicate that *Tbx1* heterozygosity impairs the acquisition speed of fimbria-dependent spatial reference memory, but not its retention or recall, or fimbria-independent visual cued memory.

**Fig. 6.**
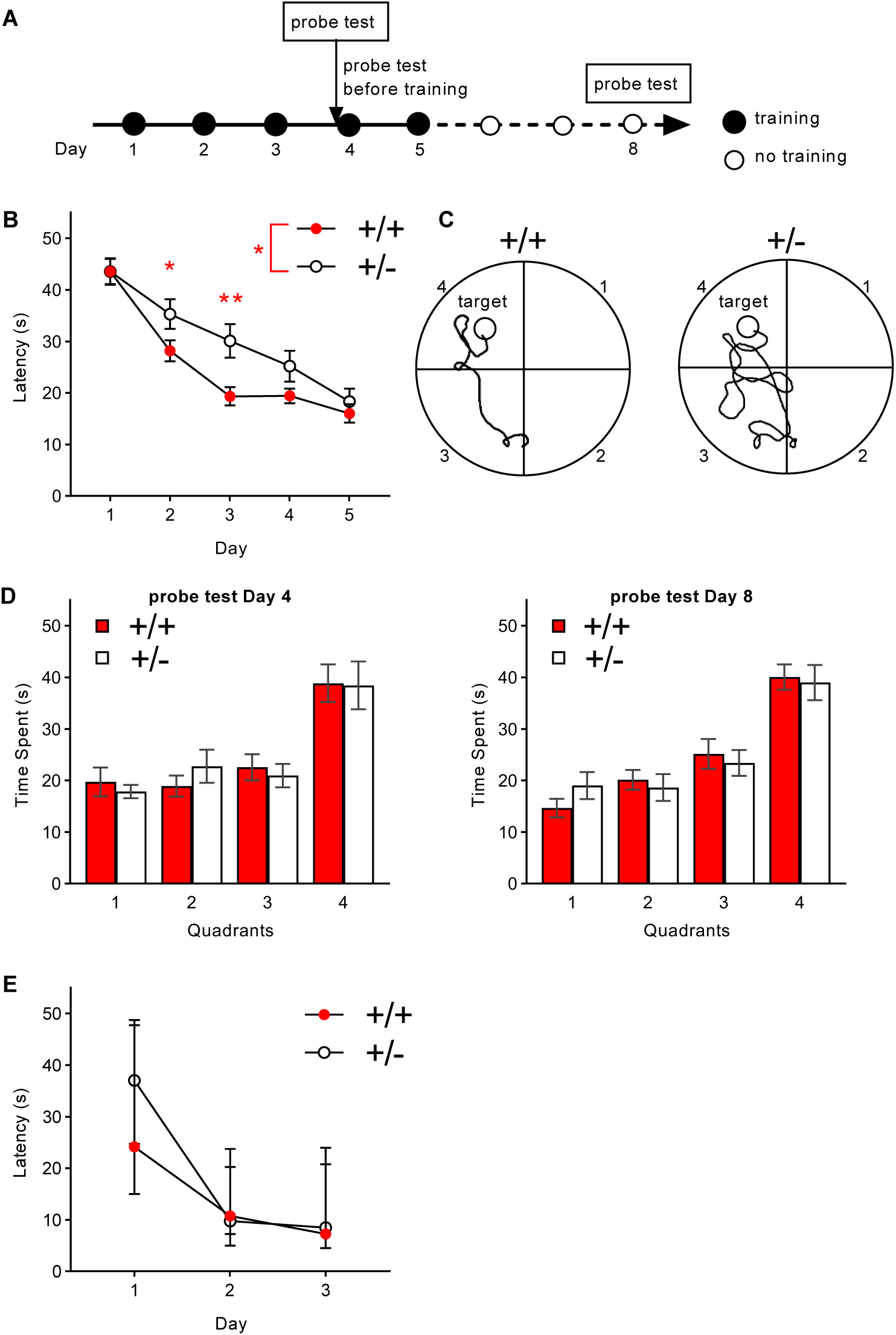
Performance in the Morris water maze test. (**A**) Experimental design. (**B**) The mean (± standard error of the mean [SEM]) escape latency in seconds (s) to the platform during acquisition is plotted against days. Compared with +/+ mice, +/- mice exhibited delayed acquisition (Genotype, F(1,26) = 4.643, p = 0.041, *[ ; Day, F(4,104) = 55.490, p < 0.001; Genotype × Day, F(4,104) = 2.329, p = 0.061). The overall genotype effect was primarily due to robust differences on Day 2 (*, p < 0.05) and Day 3 **, p < 0.01), as determined by Newman-Keuls post-hoc tests. +/+, n = 14; +/-, n = 14. (**C**) Representative swim paths of a +/+ mouse and +/- mouse on the third training day. The target quadrant included the hidden platform. (**D**) The mean (±SEM) time spent during recall probe tests before training on Day 4 (left) and Day 8 (right). Regardless of the quadrant, there were between-genotype differences on Day 4 (Genotype, F(1,26) = 5.597, p = 0.026; Quadrant, F(3,78) = 14.259, p < 0.001; Genotype × Quadrant, F(3,78) = 0.295, p = 0.829) and Day 8 (Genotype, F(1,26) = 10.207, p = 0.004; Quadrant, F(3,78) = 24.031, p < 0.001; Genotype × Quadrant, F(3,78) = 0.562, p = 0.642). The significant main effects of genotype on both days primarily resulted from the generally lower amounts of time spent in three out of the four quadrants in +/- mice (Day 4, Quadrants 1, 3, and 4; Day 8, Quadrants 2, 3, and 4). (**E**) The mean (±SEM) escape latency in the visible cue task. A separate set of mice underwent examination using this version of the Morris water maze. +/+ and +/- mice equally acquired this task (Genotype, F(1,17) = 1.861, p = 0.190; Day, F(2,34) = 52.313, p <0.001; Genotype x Day, F(2,34) = 1.229, p = 0.305) (+/+, n = 8; +/-, n = 11).

#### *Tbx1* heterozygosity slows the acquisition of discrimination and cognitive flexibility

Individuals with 22q11.2 hemizygous deletion also exhibit impairments in executive functions (*9*). Congenic mouse models of 22q11.2 hemizygosity require an increased number of trials to reach the criteria for simple discrimination and reversal learning (*56*) or extradimensional shifting (EDS) (*57*). However, the individual 22q11.2 genes contributing to impairments in executive functions remain unclear. In humans, prefrontal cortical lesions increase the number of trials required to reach the criterion of attentional set shifting; on the other hand, hippocampal lesions affect the latency for completing each trial (*58*). In rodents, orbitofrontal cortical lesions increase the number of trials required to achieve reversal of the intra- dimensional set (IDS-IV rev) (*59, 60*), although there have been no mouse studies regarding the latency for achieving attentional set shifting. *Tbx1* +/- mice lacked detectable white matter alterations in the basal forebrain or cortex (see **Fig. S2-5**) but exhibited altered myelination in the fimbria. Thus, we reasoned that *Tbx1* +/- mice may exhibit altered latency in completing attentional set shifting but may be unaffected in terms of the attentional set-shifting task requiring the prefrontal cortex (number of trials needed to reach a criterion).

There was no between-genotype difference in the number of trials required to complete each phase of attentional set shifting (**Fig. 7A**). In contrast, +/- mice were slower in completing each trial of attentional set shifting, most significantly in simple discrimination (SD) and IDS-IV rev (**Fig. 7B**).

**Figure 7.**
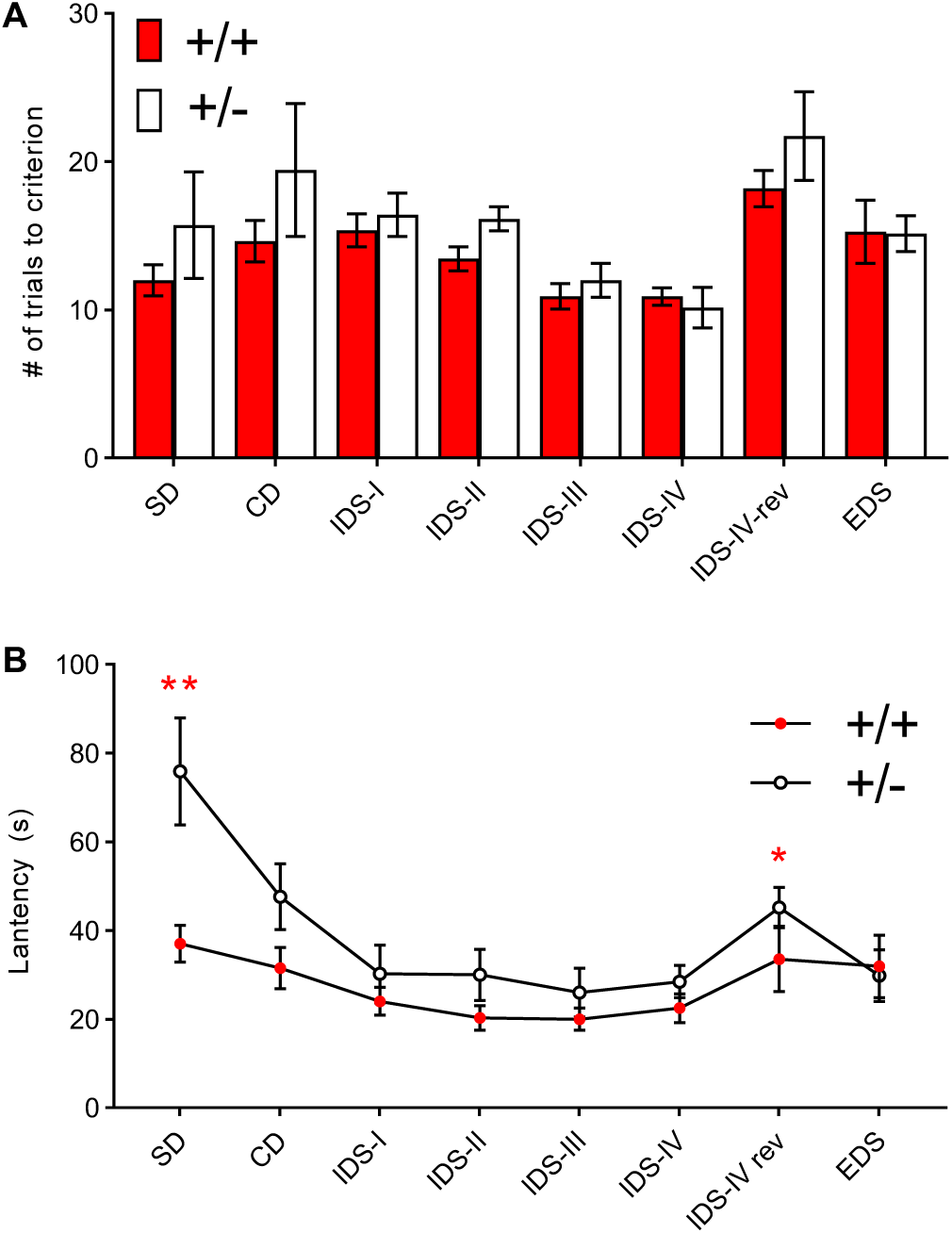
Attentional set shifting. **A**) The mean number (± standard error of the mean [SEM]) of trials required to reach the criterion (i.e., eight consecutive correct choices). Since the normality assumption was violated (p = 0.002, at IDS-IV of +/-), we performed analysis using a generalized linear mixed model. There was no between-genotype difference in the number of trials taken to reach the criterion (Genotype, F(1,16) = 1.965, p = 0.180; Genotype x Phase, F(7,112) = 0.824, p = 0.569). SD, simple discrimination; CD, compound discrimination; IDS, intra-dimensional shift; rev, reversal; EDS, extradimensional shift. (**B**) The mean latency (±SEM) to complete each trial during the first five correct choices. Since the normality assumption was violated (p = 0.001, at IDS- IV rev of +/+), a generalized linear mixed model was used for analysis. +/- mice were consistently slow in completing this task in a phase-dependent manner (Genotype, F(1,16) = 10.010, p = 0.006; Genotype x Phase, F(7, 112) = 2.566, p = 0.017). Mann- Whitney non-parametric post hoc comparisons revealed a significant between-genotype difference in latency to completing the two phases of SD (**, p < 0.01) and IDS-IV rev (*, p < 0.05). +/+ = 11, +/- = 7.

#### *Tbx1* heterozygosity has no detectable effects on olfactory responses

Consistent with the lack of detectable alterations in the white matter integrity of the neocortex, amygdala, and olfactory bulb (**Fig. S2-S5**), there was no between-genotype difference in responses or habituation to non-social and social olfactory cues (**Fig. S9**). This observation suggests that *Tbx1* heterozygosity does not exert non-specific effects on visual, olfactory, or tactile perception or on the general motivation to approach an object or odorants.

In sum, our behavioral analysis identified a highly demarcated deficit in the acquisition of fimbria-dependent cognitive tasks in Tbx1 *+/-* mice.

## DISCUSSION

The cellular, structural, or cognitive consequences of loss-of-function *TBX1* variants in humans remain unclear. A parallel analysis of structural and behavioral measures in a congenic mouse model of *Tbx1* heterozygosity indicated that *Tbx1* deficiency caused highly demarcated changes in the structural features of the brain, including reduced production of oligodendrocytes, suboptimal composition of myelination in the fimbria and corpus callosum, and loss of large myelinated axons in the fimbria and corpus callosum. These structural alterations impacted fimbria-dependent cognitive functions: *Tbx1* heterozygous mice exhibited increased latency to acquire spatial memory, simple discrimination, and reversal of intra-dimensional shift. Our findings are predictive of behavioral and structural alterations in carriers of loss-of-function *TBX1* variants in humans. Moreover, as individuals with 22q11.2 hemizygosity exhibit impairments in cognitive speed (*9, 11, 61*) as well as altered white matter integrity in the hippocampal projection fibers (*14–16, 62*), our data offer insight into the genetic and cellular substrates of these structural and behavioral alterations in carriers of 22q11.2 hemizygosity.

From a technical perspective, our combined analytical approach overcame the weaknesses associated with each technique. DTI-MRI can simultaneously screen many regions and determine the brain regions with the largest effect sizes. However, it does not identify the exact nature of altered white matter signals. Black-Gold II staining allows better resolution and detection of the reduction in net myelin density. However, gold staining was not effective in detecting the subtle effect of *Tbx1* heterozygosity on the myelination of medium axons in the corpus callosum. Although EM is labor-intensive and is not suitable for screening to identify relevant regions in the entire brain, this ultrastructural analysis revealed subtle and selective myelin alterations in large and medium axons. Therefore, a lack of detectable signal alterations in DTI-MRI or gold staining should not be considered as definitive. Moreover, an *in vitro* culture assay of oligodendrocyte precursor cells provides not only a means for evaluating gene effects on oligodendrogenesis but also a screening method for evaluating the effect of manipulating other genes and therapeutic ligands.

The observations that *Ng2* mRNA was reduced in the fimbria (see **Fig. 4A**) while markers of mature oligodendrocytes were not (see **Fig. 4A**) are seemingly difficult to reconcile. Given that the fimbria of +/- mice contained hyper-myelinated medium axons but was devoid of large myelinated axons, it is possible that the effects of these positive and negative alterations on the net amount of MBP and MOG mRNA cancel out in in the fimbria. Moreover, as myelin was selectively reduced in the anterior fimbria only (see **Fig. 2AB**), such a regionally limited effect may be difficult to detect in the whole fimbria tissue used for qRT-PCR.

The absence of large myelinated axons in the fimbria of +/- mice may be attributable to a reduced number of oligodendrocytes. Our *in vivo* data indicated that *Tbx1* heterozygosity impacts *Ng2*, a molecule required for the production of oligodendrocyte precursor cells, in the fimbria. Our *in vitro* analysis further revealed that fewer oligodendrocytes are produced from postnatal progenitor cells in +/- mice. Given their higher need for metabolic support from myelin and oligodendrocytes (*63–65*), a reduced number of oligodendrocytes may lead to degeneration of large axons. Alternatively, but not mutually exclusive, *Tbx1* heterozygosity may lead to selective inactivation of large- diameter axons and consequently reduced myelination of those axons, as oligodendrocytes tend to myelinate electrically active axons (*66*). In either case, the remaining oligodendrocytes may have instead myelinated medium axons in the fimbria, which would explain the hyper-myelination of medium axons observed in the present study.

Oligodendrocytes in the subventricular zone postnatally migrate to the fimbria and corpus callosum (*47*). In mice, Tbx1 protein is postnatally enriched in the subventricular zone (*24*). Reduced myelination of medium-diameter axons in the corpus callosum—as well as a lack of large myelinated axons in the fimbria—may have also occurred due to reduced postnatal migration of oligodendrocytes from the subventricular zone of *Tbx1* +/- mice (see **Fig. 3EF**). It remains unclear how local oligodendrocyte precursor cells in the fimbria and oligodendrocytes postnatally provided from the subventricular zone contribute to the myelination of axons of different sizes. There is a need for further studies to explore the molecular mechanisms underlying the role of *Tbx1* in myelin composition within the fimbria and corpus callosum.

Robust structural alterations in the fimbria exerted effects on the acquisition speed of spatial memory in the Morris water maze and cognitive flexibility in the attentional set shifting task. The first day of the Morris water maze reflects chance-level performance since the mice have not acquired a spatial map for the location of the platform. From day 2 onward, the speed of mastering the spatial map is reflected by the latency to reach the platform, with +/- mice exhibiting a significant delay. When mice encounter the first discrimination task or reversal of intra-dimensional shift task, they are likely to face difficulty. In the present study, +/+ mice exhibited longer latencies to complete those phases than other phases (see **Fig 7B**, SD and IDS-IV rev). It is noteworthy that +/- mice exhibited the most significant delays in completing those phases, suggesting that *Tbx1* deficiency impairs the ability to quickly master cognitively difficult tasks.

Our single-gene analysis provides a valid first step for deconstruction and reconstruction of the mechanistic composition of CNV-encoded genes in terms of their association with specific behavioral and structural dimensions. We previously reported that gene-dose alterations of *Tbx1* impair reciprocal social interaction (*21, 24*), social communication (*27, 30*), and working memory (*24, 28*). The present findings further demonstrate the effects of *Tbx1* heterozygosity on the myelin composition of the fimbria and its cognitive functions. However, we cannot exclude the possibility that deficiency of other 22q11.2 driver genes also contributes to similar and other cognitive deficits (*29*). Moreover, it is possible that other cellular mechanisms exist for social interaction and communication deficits of *Tbx1* heterozygosity. Since 22q11.2 CNVs also include genes without an apparent role in any dimension (i.e., non-contributory genes) (*20, 22, 29, 67, 68*), there is a need to comprehensively investigate each encoded single gene to elucidate the mechanisms underlying the effects of this CNV on behavioral dimensions. It should be cautioned, however, that a genotype may not impact a dimension as a unit and instead impact variables within a dimension (*69*)

The structural alterations and cognitive deficits observed in the present study are not unique to *Tbx1* heterozygosity or 22q11.2 CNVs. Lower FA values have been reported in the fimbria/fornix of individuals with idiopathic ASD (*70*) and schizophrenia (*71*). Slow processing speed in individuals with idiopathic ASD is correlated with low FA values, but not with MD, RD, or AD values, in the whole brain (*72*). Individuals with idiopathic ASD also exhibit impairments in difficult cognitive tasks (*73*). A selective loss of extra-large myelinated axons has been observed in the brains of humans with ASD (*74*). Moreover, patients with idiopathic schizophrenia exhibit impaired processing speed across numerous cognitive dimensions, including attention, memory, spatial processing, emotional identification, and sensorimotor capacity (*75, 76*). Previous studies have also reported that other oligodendrocyte-related genes are dysregulated in brain samples from individuals with ASD and genetic mouse models for ASD (*77–81*). Taken together, our findings open a new window for investigating the potential substrates of altered cognitive speed in carriers of *TBX1* SNVs, 22q11.2 and other CNVs, and in idiopathic cases of ASD and schizophrenia.

### Limitations

We screened for specific brain regions using DTI-MRI imaging. However, our resolution (150 μm isotropic voxel) may not have been sufficient for detecting subtle alterations. Although DTI-MRI analysis revealed significant differences in the FA values for the fimbria only, EM analysis revealed less myelination exclusively in medium-sized axons in the corpus callosum of +/- mice as well. If the gene deficiency affects a specific structural set or axons with a certain diameter, it would be difficult to detect such subtle effects using DTI-MRI. Therefore, the finding regarding the absence of detectable alterations based on the DTI-MRI analysis of other regions should be interpreted cautiously.

We interchangeably used male and female mice for various analyses, as individuals with 22q11.2 hemizygosity do not exhibit a sex bias for schizophrenia or ASD diagnosis (*82*) or for various cognitive capacities, including set-shifting, memory, and processing speed (*9, 11*). The number of currently identified *TBX1* loss-of-function mutations is too small to determine a sex bias, however. Thus, there is a need for further research to determine the precise impact of sex on various phenotypes.

## MATERIALS AND METHODS

### Experimental Design

This study was designed to determine the structural and cellular bases underlying specific cognitive functions affected by *Tbx1* heterozygosity in a congenic mouse model. Specifically, we screened the most robust microstructural alterations using DTI-MRI, histologically validated the findings through gold staining, identified ultra-structural bases using EM, and determined the *in vitro* oligodendrocyte production capacity. After demonstrating that *Tbx1* heterozygosity alters myelin composition in the fimbria, we evaluated fimbria-dependent and fimbria-independent cognitive functions using the Morris water maze, attentional set shifting, and olfactory responses and habituation.

### Mice

The protocols for animal handling and use were approved by the Animal Care and Use Committee of the Albert Einstein College of Medicine, University of Texas Health Science Center at San Antonio and Tohoku University in accordance with National Institutes of Health (NIH) guidelines.

*Tbx1*^+/-^ mice This mouse model was a congenic strain with a C57BL/6J background. The original non-congenic *Tbx1*^+/-^ mouse was backcrossed onto C57BL/6J inbred mice for >10 generations to control for biased genetic backgrounds (*83*). Given that there are no sex biases in the prevalence of schizophrenia or ASD (*82*), set-shifting, spatial working memory, spatial planning, processing speed, or other cognitive domains (*9, 11*) among carriers of 22q11.2 hemizygosity, we used either male or female mice for the various analyses.

We determined genotypes of mice using three primers: forward TTGGTGACGATCATCTCGGT and reverse ATGATCTCCGCCGTGTCTAG to detect the +/+ genotype, as well as an additional reverse AGGTCCCTCGAAGAGGTTCA to detect the +/- genotype.

#### Sample preparation

We performed *ex vivo* MR scanning to achieve a high resolution and high signal-to-noise ratio since it allows a long scan time and involves the use of a contrast agent. In accordance with standard procedures (*84*), 4-month old female mice were anesthetized using pentobarbital (60 mg/kg, i.p.) and transcardially perfused using 30 mL of 0.01 M phosphate-buffered saline (PBS) that contained 2 mM of ProHance (Bracco-Eisai Co., Ltd, Tokyo, Japan) and 1 μL/mL heparin (1,000 USP units/mL), followed by 30 mL of 4% paraformaldehyde (PFA; Wako, Tokyo, Japan) containing 2 mM ProHance. The head was decapitated, following which the skin, lower jaw, ears, and cartilaginous nose tip were removed. The skull structure containing the brain tissue was post-fixed in fixative (4% PFA and 2 mM ProHance) overnight at 4°C. Subsequently, it was transferred to buffer (0.01 M PBS, 0.02% sodium azide, and 2 mM ProHance) at 4°C overnight. Next, the brain tissues were placed in fresh buffer (0.01 M PBS, 0.02% sodium azide + 2 mM ProHance). Immediately before scanning, we immersed *ex vivo* mouse brains in Fomblin (Sigma-Aldrich, St Louis, MO), which is a perfluorocarbon that reduces susceptibility artifacts at the interface and limits intra-scanning sample dehydration.

#### MRI acquisition

MRI data were acquired using a 7.0-T PharmaScan 70/16 system with a 23-mm diameter birdcage Tx/Rx coil specifically designed for the mouse brain (Bruker Biospin, Ettlingen, Germany) using standard operational software (Paravision 6.0.1). We acquired triplot images to ensure proper sample positioning with respect to the magnet isocenter. Shim gradients were adjusted using the MAPSHIM protocol with an ellipsoid reference volume covering the whole brain. We obtained diffusion-weighted images using a standard spin-echo 2D pulse sequence using the following parameters: repetition time = 4,158 ms, echo time = 42 ms, field of view = 15 × 12 mm^2^, matrix size = 100 × 80, in-plane resolution = 0.15 × 0.15 mm^2^, number of slices = 50, slice thickness = 0.3 mm, diffusion gradient duration = 6 ms, diffusion gradient separation = 30 ms, b-value = 2,000 s/mm^2^, number of diffusion directions = 30, number of b0 images = 1, effective spectral bandwidth = 30 kHz, fat suppression = on, and number of averages = 10. The diffusion- weighted images was acquired at 22^0^C-26^0^C for 22 h per mouse.

#### Diffusion-weighted image analysis

Acquired images were processed using the Advanced Normalization Tools (http://stnava.github.io/ANTs/) and FMRIB Software Library (FSL) software packages (https://fsl.fmrib.ox.ac.uk/fsl/fslwiki). The procedure for image processing was as follows: (i) Image reconstruction was performed using Paravision software and converted to the NIfTI format using “DSI Studio” software (http://dsi-studio.labsolver.org/); (ii) eddy-current induced distortions were corrected using the eddy_correct tool of FSL; (iii) individual reference b0 images were manually skull-stripped using ITK-SNAP software (http://www.itksnap.org); (iv) other subject b0 images were registered to the reference image and skull-stripped; (v) scalar images were reconstructed using the DTIFIT tool of FSL; (vi) b0 and scalar images were manually rotated and translated to ensure that the coordinate origins occupied the anterior commissure midpoint to roughly match the standard reference space; (vii) b0 and scalar images were resampled onto an 0.15-mm isotropic voxel; (viii) the Minimum Deformation Template (MDT) space was constructed using all subject b0 and scalar images, including FA, AD, MD, and RD images; (ix) b0 and scalar images were warped to the MDT space; (x) the mean b0 image was computed, manually skull-stripped, and registered to the atlas image; and (xi) the mean FA, AD, MD, and RD values in each structure were computed and statistically analyzed.

### Black-Gold II staining

Two-to three-month-old female mice were perfused with saline and 4% paraformaldehyde as per the standard protocol (*28*). We mounted a pair of free-floating 40-μm thick coronal sections from a +/+ mouse and a +/- littermate as the upper and lower rows, respectively, on the same slides (3-4 section pairs per slide) to control for cross-slide staining variations. Care was taken to mount a section pair with similar coordinates from +/+ and +/- mice on the upper and lower rows of a slide.

The degree of myelination was examined using Black-Gold II staining (*85*). Black-Gold II is an aurohalophosphate complex that directly stains myelin within the CNS. Black-Gold II and sodium thiosulfate solution (AG105, Millipore, Temecula, CA) were heated to 60°C. Slide-mounted sections were rehydrated in filtered water, transferred to pre- warmed Black-Gold II solution, and incubated at 60°C for >12 min. Subsequently, the sections were rinsed in filtered water twice for 2 min each, transferred to sodium thiosulfate solution, and incubated for 3 min at 60°C. Finally, the sections were rinsed three times in filtered water for 2 min each and cover-slipped.

We semi-quantified gold-staining within the fimbria and corpus callosum using a Keyence microscope and its controller (BZ-X810 and BZ-X800E). Under a light microscope, staining blocked light penetration through the sections and registered as less bright. This property was employed for semi-quantitative analysis.

The fimbria and corpus callosum regions were delineated as targets (see **Fig. S6**). We divided the fimbria and corpus callosum at Bregma –1.30 mm, the anterior and posterior of which were defined as anterior and posterior areas of the two regions for analysis.

The Keyence software yields brightness (B) values as integration values within a range of threshold values from 0 to 255. The threshold value acts as a filter and determines the level of light that is allowed to penetrate through a section. We observed that sampling pixels gradually saturated areas where tissues exist up to a threshold unit value of 137, above which pixels started to appear non-specifically in areas devoid of tissue (e.g., blood vessels and between-tissue gaps). Thus, signals are maximally detected without false positive signals at this threshold value. This threshold was consistently used in the analysis of staining signals. Since B represents the sum of all integration values within the delineated area, it is affected by the size of the area. Because the target area size varied from section to section, we computed B per area (A) unit of the target (t) region (i.e., tB/A) in the fimbria and corpus callosum (**S-Table 1, Step 1**).

Although we minimized slide-to-slide variations in staining intensity by dipping a set of slides in the same Black-Gold II solution, there was still variation. This was observed as varying non-specific baseline staining across sections. To correct for this variation, we adjusted the tB/A value based on the degree of non-specific staining. We chose a 250 μm × 250 μm cortical area above the target fimbria and corpus callosum, where gold- labeling was negligible. We defined it as a negative control (nc) area where B/A values represent non-specific staining (**S-Table 1, Step 2**). The threshold unit for genuine tissue signals was 255 in the cortex, above which signals started to appear in areas with no tissue. We next chose a section with the maximum negative control B/A (max ncB/A) value and converted all ncB/A values to ratios (R=(max ncB/A)/(ncB/A), **S-Table 1, Step 3**).

Next, the tB/A value was multiplied by the R value, such that an under-estimated brightness signal due to non-specific staining (i.e., low B/A value) was rectified proportionally to the relative degree of non-specific staining (tB/A^adj^=(tB/A)*R, **S-Table 1, Step 4**).

As staining intensity is inversely proportional to the tB/A^adj^ value, greater gold staining indicates that less light penetrates a section. The inverse value of tB/A^adj^ was calculated (1/(tB/A^adj^), **S-Table 1, Step 5**) and multiplied by 10^3^ to express values above the decimal point.

### EM analyses

Two- to three-month-old male mice were anesthetized using 4-5% isoflurane in a chamber, and anesthesia was maintained with 2.0-3.5% isoflurane using a nose-cone vaporizer. The animals were intracardially perfused with 100 mL of 0.9% physiological saline followed by approximately 250 mL of freshly prepared 0.1 M sodium cacodylate buffer (pH 7.4; Electron Microscopy Sciences cat #11653), which contained 2.5% glutaraldehyde (Electron Microscopy Sciences cat #16320) and 2.5% PFA (Electron Microscopy Sciences cat #19202). Next, the brains were split into two hemispheres and post-fixed in fixative at 4°C for 2 weeks. Samples from the target areas (fimbria and corpus callosum) were obtained using a vibratome and placed in 0.1 M sodium cacodylate buffer overnight. The tissues were then rinsed three times for 10 min each in 0.1 M cacodylate buffer to remove aldehydes, following which they were placed in a mixture (500 μL) of 2% OsO_4_ (Electron Microscopy Sciences, cat#19150) and 0.1 M sodium cacodylate buffer for 1 h. The tissue samples were agitated and shaken, rinsed (3 x 5 min 0.1 M Na cacodylate), and dehydrated twice in a series of ice-cold ethanol solutions for 5 min each (30% ethanol; 50% ethanol; 70% ethanol; 90% ethanol; 95% ethanol) and three times in 100% ethanol for 10 min. Next, the tissues were rinsed twice in propylene oxide for 30 min each (Polysciences, Inc., cat# 00236–1). This was followed by incubation on a mixer at room temperature overnight in an approximately 1 mL mixture of 1 part propylene oxide and 1 part Polybed resin solution (Poly/Bed® 812 Embedding Media, Polysciences, Inc., cat# 08791–500; Dodecenylsuccinic anhydride (DDSA, Polysciences Inc., cat# 00563–450), nadic methyl anhydride (NMA, Polysciences Inc., cat# 00886–500), and 2,4,6-Tris-(dimethylaminomethyl)phenol (DMP- 30, Polysciences, Inc., cat# 00553–100). On the next day, the Polybed resin/propylene oxide solution was removed, and the tissues were incubated for 24 h in 100% Polybed solution on a mixer at room temperature. Tissues were removed from the Polybed resin and placed in a mold, following which fresh polyresin was added. After the resulting bubbles had disappeared, the tissues in the mold were incubated at 55°C for 36 h. Subsequently, they were processed at the Electron Microscopy Laboratory of the UT Health Science Center in San Antonio using the in-house procedure (*86*). The tissues were cut at 1 µm and stained using 0.1% toluidine blue/0, 0.1% methylene blue/0, and 0.1% azure II in 1% sodium borate buffer. Next, 100-nm thick sections were cut and collected on 300 hexagonal mesh copper grids (Electron Microscopy Sciences, cat # T300H-Cu). In each set of five grids, three were stained, and two were left unstained. Staining was performed using 5% uranyl acetate in 50% methanol and Reynold’s lead citrate (*87*). We measured the diameters of myelinated axons and their axon portions.

### qRT-PCR

We used 2-3 month-old female *Tbx1* +/+ and +/- littermates. Total RNA was extracted from brain regions of adult mice using an RNeasy Plus Mini Kit (Cat#74134, Qiagen, Germantown, USA), in accordance with the manufacturer’s instructions. cDNA was synthesized from total RNA using SuperScript IV VILO master mix (Cat# 11766050, Invitrogen, Carlsbad, USA). Quantitative PCR reactions were performed in triplicate on QuantStudio 6 Flex Real-Time PCR Systems (Cat#4485694, Applied Biosystems, Waltham, USA) using the TaqMan Fast Advanced Master Mix (Cat#4444963, Applied Biosystems, Waltham, USA). The Taqman probes are listed in the Supplementary Material (**Table S3**). Data were analyzed using the ΔΔCt method and normalized to the reference gene Cyc1.

### *In vitro* analysis of oligodendrocytes

We used P21 +/+ (n =9) and +/- mice (n = 5) chosen from five litters. Progenitor cells were isolated from the lateral ventricular walls of both hemispheres. Two 1mm slices were taken from each of both hemispheres, and tissues that include the subventricular zone were dissected. Each culture was prepared using tissue from a single mouse. The tissues were dissociated using a Neural Tissue Dissociation Kit (P) (130-092-628, Miltenyi Biotech GmbH, Germany). We did not purify neural progenitor cells with antibodies; thus, our cells contained different types of proliferating progenitor cells that generate neurons and oligodendrocytes. The cells were cultured in a medium (DMEM/F12 [11320–033, Gibco, CA, USA]) supplemented with N2 (17502048, Gibco), B27 (17504044, Gibco), epidermal growth factor (EGF) (20 ng/ml) (AF100-15, Peprotech, NJ, USA), and fibroblast growth factor 2 (FGF2) (10 ng/ml) (100-10B, Peprotech). After two to three passages, the cells were dissociated from the spheres and seeded on a Matrigel (356234, BD Biosciences, Bedford MA, USA)-coated slide chamber (154534, Nunc, NY, USA). To promote differentiation, the cells were cultured for 4 days in medium supplemented with 5% fetal calf serum. Next, the cells were fixed using 4% PFA for 15 min and processed for immunofluorescence staining, using a purified mouse monoclonal O-4 antibody (1:50, MAB345, Millipore, MA, USA) for 12 h at 4°C after blocking for 30 min at room temperature with 5% donkey serum (S30-100ml, Millipore, MA, USA). Subsequently, the cells were incubated for 30 min at room temperature with Goat anti-Mouse IgM (Heavy Chain) Secondary Antibody, Alexa Fluor 647 (1:1000, A21238, Molecular Probes, OR USA). Nucleus staining was performed using 4’,6-diamidino-2-phenylindole (DAPI) (3 mM, D3571, Molecular Probes). Cells were counted from four randomly selected fields per culture under a confocal microscope (TCS SP8, Leica, Germany), and the average score was obtained.

### Behavioral analysis

The mice were tested during the light phase between 10 AM and 5 PM.

#### Morris water maze

Separate groups of 2-month-old male mice were used for the hidden and visible platform versions of the Morris water maze test. The water tank (103 cm in diameter; ∼914 lx) contained white Prang® (Dixon Ticonderoga®) Ready-to-Use Paint (Item #: 738062, Model #: 21609/21949, Staples) mixed in water (24 ± 2°C). A circular platform (10 cm in diameter) was submerged 1 cm below the surface in the middle of one quadrant. Cues were placed on the wall 40 cm from the tank edge. The water was changed after testing on days 3 and 5.

The hidden platform version involved 10 sessions conducted over 5 days (two daily sessions at intervals of 2–4 hours). Each session included four 60-s trials conducted at 15-min intervals. The platform location remained constant (Quadrant 4); however, the entry points were semi-randomly changed across the trials. Before the fourth day of the hidden platform training, we performed a 60-s probe trial for which the platform was removed. The entry point for the probe trials was the quadrant opposite to the target quadrant. An additional probe trial was conducted 72 h after the fifth day of hidden- platform training.

The cued platform version involved six sessions conducted over 3 days (two daily sessions at intervals of 2–4 h with each session having two 60-s trials at 15-min intervals). The platform was marked using a flag placed above the water surface and visible to the mice. The platform locations were randomly assigned to each trial. The mice were placed in the maze from four equally spaced points along the pool perimeter, and the entry-point sequence was randomly chosen. For each placement, the animals were placed facing the sidewalls. The sequence of four start positions (north, south, east, and west) varied across the trials.

In both the hidden and cued platform versions, all animals were allowed to remain on the platform for 30 s. In case they did not reach the platform during the 60-s test, the experimenter placed and left the animal on the platform for 30 s. During the 15-min inter- trial interval, the mouse was dried using a paper towel and placed in an empty cage stuffed with a dry paper towel.

#### Attentional set shifting

This test was performed using a procedure optimized for mice (*59*), with a slight modification. Two- to three-month-old male mice were individually housed and food-deprived to reduce the bodyweight to 85% of the *ad libitum* feeding weight, and this bodyweight was maintained throughout the testing period.

The mice were taken to the test room 1 h before the start of the training session. A single bowl containing 1/2 of a Honey Nut Cheerio buried in one medium stimulus sprinkled with an odor stimulus was placed in the home cage. This training used all possible combinations of exemplars of both dimensions (i.e., odor and medium) for subsequent use in the eight phases of attentional set shifting. The mice completed four daily trials, each involving a unique combination of medium and odor stimuli. The bowl was immediately removed from the home cage after the mouse had dug up the food pellet and eaten it. Each trial lasted approximately 1–2 min. After completing the daily training trials, the mouse was placed in a new home cage with fresh bedding.

Next, we conducted a one-day habituation session in the attentional set-shifting apparatus (outer dimensions: height (H), 15 cm x width (W), 19.2 cm x length (L), 49.2 cm; inner dimensions: H, 14.4 cm x W, 18.3 cm x L, 48.3 cm; ∼914 lux). The apparatus was divided into two goal compartments (W, 9 cm x L, 14 cm each) and one start compartment (W, 18.3 cm x L, 33.9 cm) using 4.8-mm-thick walls. The mouse explored the apparatus arena, which included a plastic weigh boat containing water in the start compartment. The two goal compartments lacked bowls. After 3 minutes, the partition door was placed to confine the mouse to the start compartment. After another 3 minutes, the door was removed to allow the mouse to freely explore all three compartments. The two 3-min sessions were repeated five times.

On the next day, training began in the attentional set-shifting session apparatus. The water tray remained in the starting compartment during testing and initial re-training. The two bowls in the goal compartments contained two medium stimuli (e.g., alpha dri and paper chips) without odor stimuli; moreover, they were both baited using food. Both media were used for the subsequent SD sessions. The partition door was placed to confine the mouse to the start compartment. Care was taken to remove the door when the mouse was not sniffing or facing it. Initially, the mice underwent four re-training trials to retrieve the food from the bowl. The positions of both medium-containing bowls were randomized in each trial. We placed an eighth of a Honey Nut Cheerio on top of (trial 1), half-buried within (trial 2), slightly covered by (trial 3), and completely buried within (trial 4) the media. Each trial ended when the mouse had eaten food from both bowls.

Subsequently, the mice underwent a series of discrimination tests. Initially, each mouse was placed in the start compartment with a partition door. The two goal compartments contained one baited bowl (an eighth of a Cheerio piece completely buried in the medium) and one un-baited bowl; moreover, the position of the baited bowl was randomized across the trials. The partition door was then lifted. Each trial ended when the mouse had made a correct choice and had eaten the reward. If the mouse dug into the un-baited bowl, it was removed after the mouse had spontaneously left the un-baited compartment. A time-out was given if the mouse did not dig in any bowl for 3 min, which involved removal of the bowl from the test arena and subsequent resumption of the trial using a different medium/odor pair. In case of three consecutive time-outs, the testing was ended and resumed the next day. Each of the eight phases ended when the mouse had made eight consecutive correct choices or after 50 trials per day, whichever came first. If a mouse made eight consecutive correct choices within 50 trials, a new phase was administered the next day. After each test trial and when changing mice, the arena and bowls were wiped using 70% ethanol.

The attentional set-shifting phases were as follows (**Tables S4** and **S5**). For the SD phase, there were two choices for the two relevant dimensions. The compound discrimination (CD) phase was similar to the SD phase, except that a new correct compound (O1&M1 and O1&M2) was added. The IDS IV phase involved CD using two novel exemplars from relevant and irrelevant dimensions for each IDS with the same relevance. The IDS IV rev phase involved the same exemplar set as the IDS IV phase, except that the correct choice within the relevant dimension was reversed. The extradimensional shifting (EDS) phase involved novel CD, except that the correct choice was an exemplar of the previously irrelevant dimension up to IDS-IV rev. The order of discrimination and exemplars was similar for all mice. The exemplar choice and correct bowl position were pre-determined using a random number table.

The standard mouse SD procedure uses O1 plus M1 and O2 plus M1 (*59*). Our modified SD procedure used a combination of two dimensions (O1 plus M1 as the correct discriminants and O2 plus M2 as the incorrect discriminants). Our pilot study indicated that, compared with +/+ mice, +/- mice exhibited a longer latency to complete this modified task.

We determined the number of trials taken to reach eight consecutive correct choices and the latency to complete a trial from the trial start to the time point when the mice began eating the food pellet.

#### Olfactory responses to social and non-social cues

This test was conducted in a test cage (L, 28.5 cm × W, 17.5 cm × H, 12.5 cm) that had been divided into a 19.5 cm-long compartment and a 9 cm-long compartment using a partition wall with a 5 cm (H) × 5 cm (W) opening; ∼430 lux. The test was conducted as previously described (*24*), with slight modifications. First, 2-month-old male mice were habituated to the apparatus for 15 min. A filter paper scented with a test odor was placed in a 1-ml Eppendorf tube containing small holes in the cap. Odors were sequentially tested as follows: water, almond, banana, urine from one non-littermate C57BL/6J male (NL1), urine from another non-littermate C57BL/6J male (NL2), urine from the first C57BL/6N mouse (NL1), urine from a non- littermate male +/- mouse (HT), urine from the dam (rm), urine from another litter’s mother (am), and urine from a non-littermate virgin female C57BL/6J mouse (v). We measured sniffing of the tube containing odorant-soaked filter paper during the 2-min trials. The mice underwent three 2-min trials for each odorant with an inter-trial interval of approximately 10 s; moreover, there was a 10-s interval between the three-trial session of one odorant and that of another. Urine was collected before testing and frozen at –20 °C until the test day. An Eppendorf tube with seven holes (one in the middle and six surrounding) in the cap was used for each trial. The tube was attached to the cage wall using Velcro. The filter paper (Whatman, #3698-325, Maidstone England) was soaked in 10 µl of each odorant. During habituation, we placed dry filter paper in the tube.

### Statistical analysis

We used GraphPad Prism 8.3.0 (GraphPad Software, San Diego, CA) and IBM SPSS Statistics 26.0.0.0, IBM, Armonk, NY). Among-group and between-group comparisons of the data were performed using analyses of variance and Student’s t-test, respectively. Normality and variance homogeneity of the data were evaluated using the Shapiro-Wilk test and Levene’s homogeneity of variance test, respectively. In case either assumption was violated, data were analyzed using a generalized linear mixed model or Mann-Whitney U-tests and Wilcoxon non-parametric tests for unpaired and paired data, respectively. The number of cases was analyzed using the χ^2^ test. The minimum significance level was set at 5%. In case multiple tests were applied for a data set, the significance level was adjusted using the Benjamini-Hochberg correction, with a false discovery rate of 5%.

## Supporting information

Hiroi Supplemental materials

## Acknowledgments

**General**: We thank Dr. Bernice Morrow for providing the original breeders of *Tbx1*^+/-^ mice.

**Funding**: This study was partly supported by the National Institutes of Health (R01MH099660, R01DC015776, R21HD053114). The content is solely the responsibility of the authors and does not necessarily represent the official views of the National Institutes of Health.

**Author Contributions**: T. Hiramoto and N. Hiroi designed the study and analyzed all the data. A. Sumiyoshi, R. Ryoke, H. Nonaka, and R. Kawashima designed and performed the DTI-MRI study and analyzed the data. T. Yamauchi performed gold staining immunohistochemistry and qRT-PCR and analyzed the data. G. Kang maintained and genotyped the mouse colony of *Tbx1*^+/-^ mice and performed behavioral studies, except for the Morris water maze and attentional set shifting tasks. T. Hiramoto performed the Morris water maze test and analyzed the data. T. Hiramoto., S. Enomoto, and T. Izumi performed attentional set shifting and analyzed the data. K. Tanigaki performed *in vitro* cell cultures of oligodendrocytes and analyzed the data. T. Hiramoto, A. Sumiyoshi, R. Kato, T. Yamauchi, G. Kang, K. Tanigaki, and N. Hiroi wrote the manuscript.

**Competing interests**: None

**Data and material availability**: All data are available upon request. Mice are available through the Material Transfer Agreement.

## Notes

### Competing Interest Statement

The authors have declared no competing interest.

### Summary of Updates

The original Fig 4 is split into Fig 4 and Fig 5. A new set of images are used for Fig 5.

