## Supplementary material for "*Tbx1*, a 22q11.2-encoded gene, is a link between alterations in fimbria myelination and cognitive speed in mice": Hiroi Supplemental materials

**This PDF file includes:**

Figs. S1 to S9

Tables S1 to S5

Reference

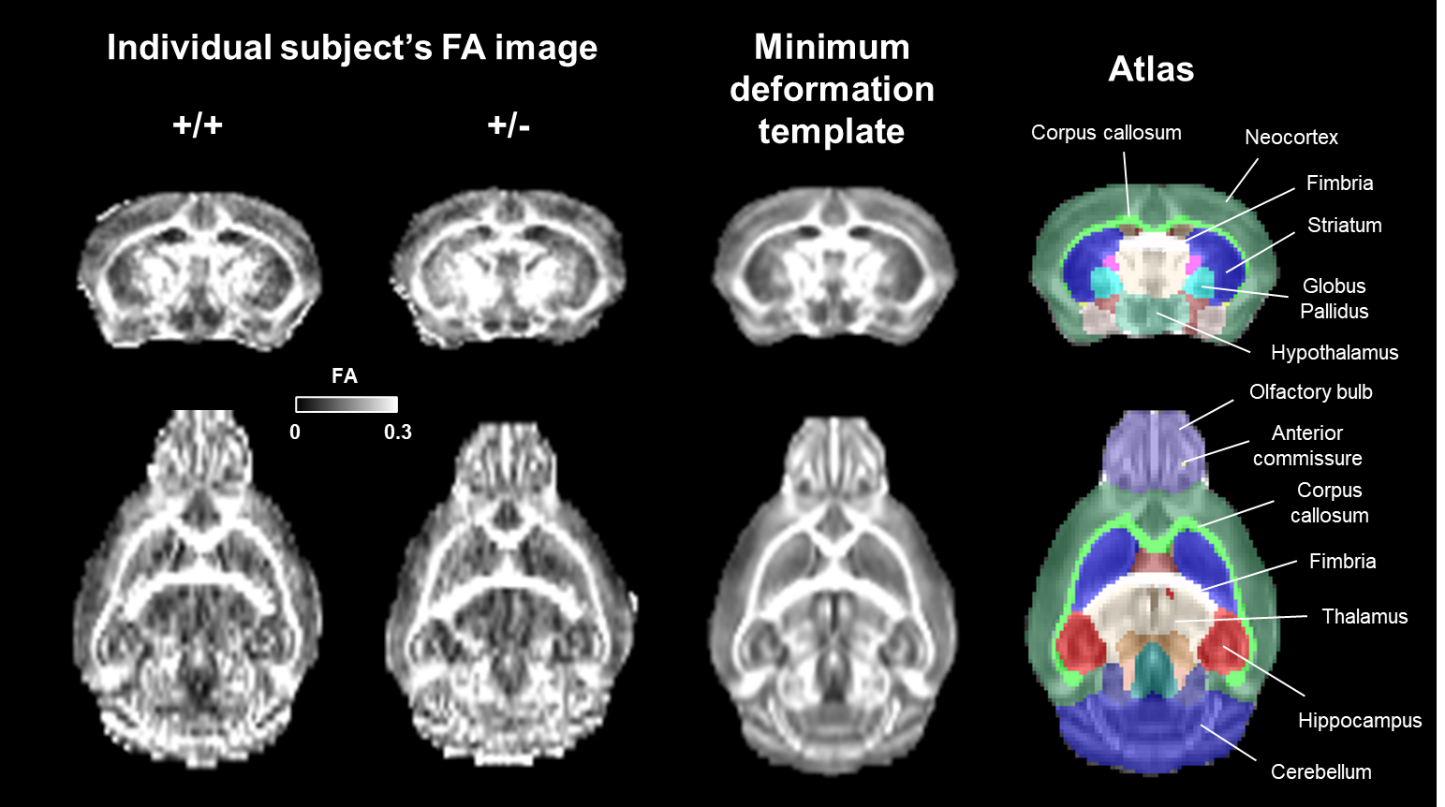
**Figure S1.** High-resolution *ex vivo* fractional anisotropy (FA) image (150-μm isotropic voxel). The minimum deformation template was constructed from all b0 and scalar images (N = 17) and nonlinearly registered to the mouse atlas space (*1*). Each anatomical structure was used for statistical analyses.

**A**
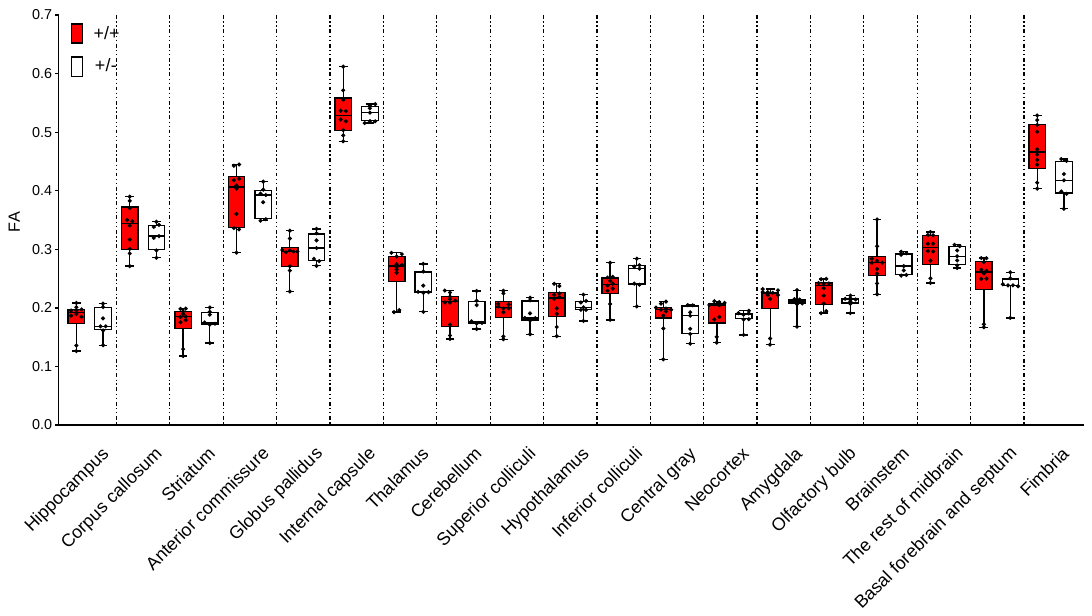

**B**
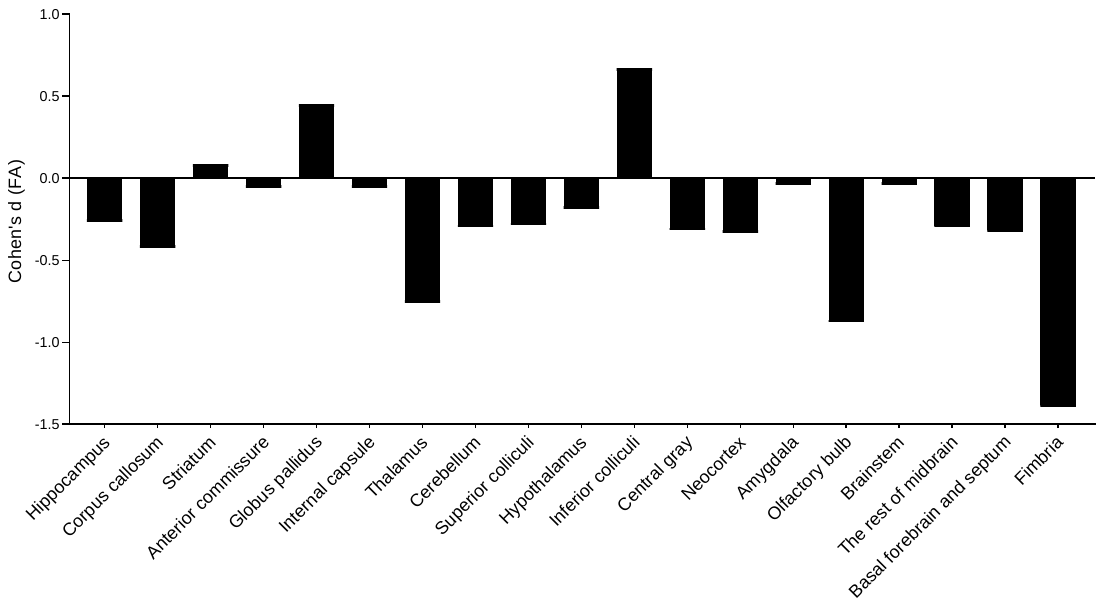

**Figure S2.** (**A**) There were between-genotype differences in fractional anisotropy (FA) values in some regions (Genotype x Region, F(18, 270) = 2.795, p < 0.001). (**B**) Effect sizes of the FA differences between +/+ and +/- mice, as determined by Cohen’s d values. +/+, n = 10; +/-, n = 7.

| **A**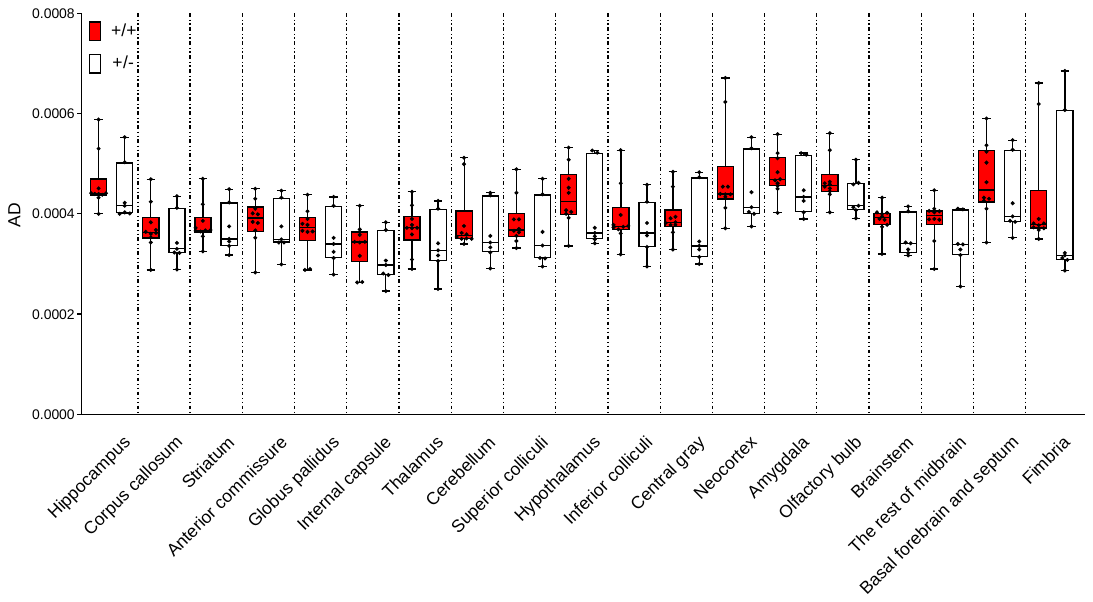  **B**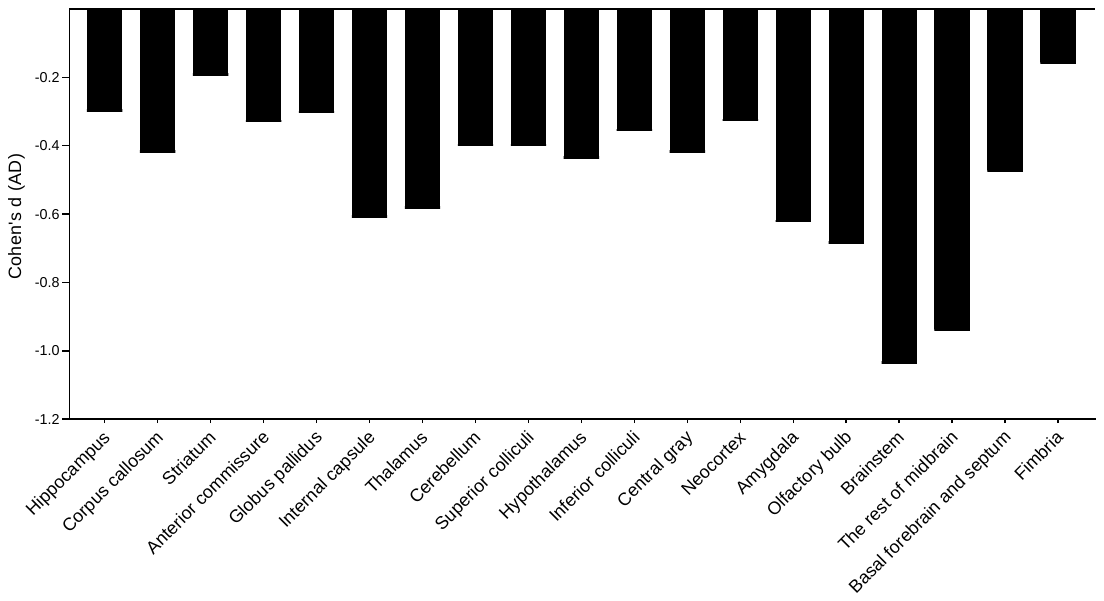 |
| --- |

**Figure S3. (A)**. Mann-Whitney U-tests revealed no between-genotype differences in axial diffusivity (AD) values in any of the regions examined (all p > 0.05). (**B)**. Effect sizes of the AD differences between +/+ and +/- mice, as determined by Cohen’s d values.

**A**
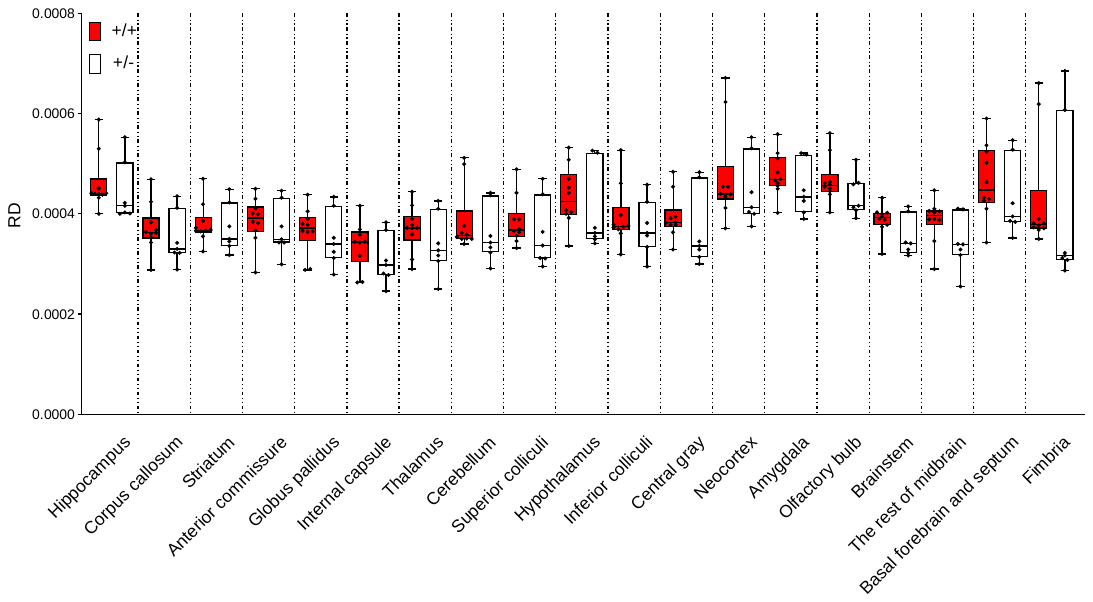

**B**
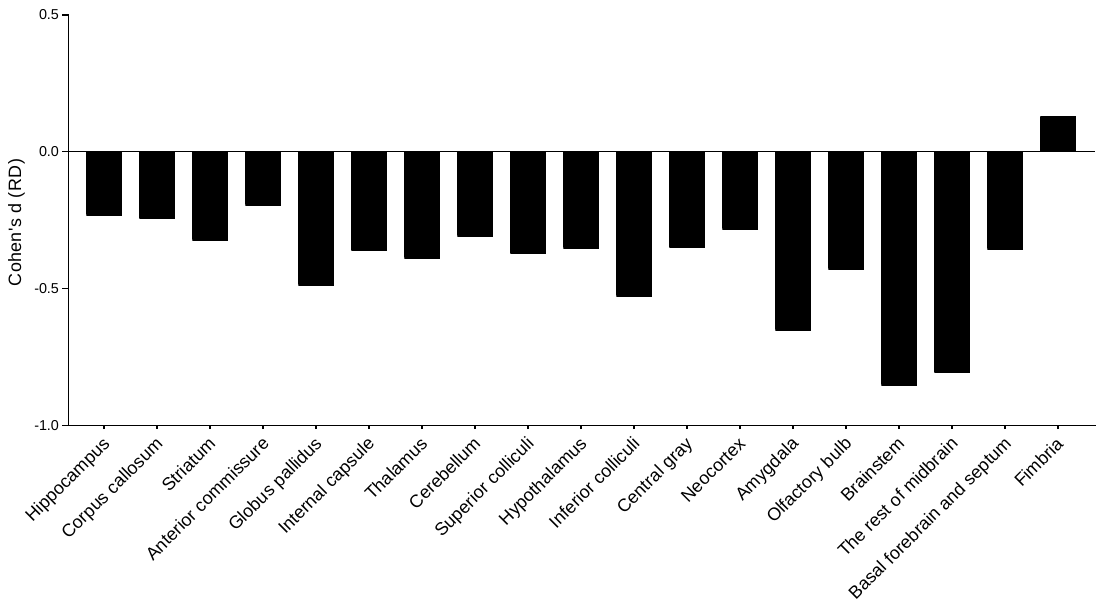

**Figure S4**. **(A)**. Mann-Whitney U-tests revealed no between-genotype differences in radial diffusivity (RD) values in any of the regions examined (all p > 0.05). (**B)**. Effect sizes of the RD differences between +/+ and +/- mice, as determined by Cohen’s d values.

**A**
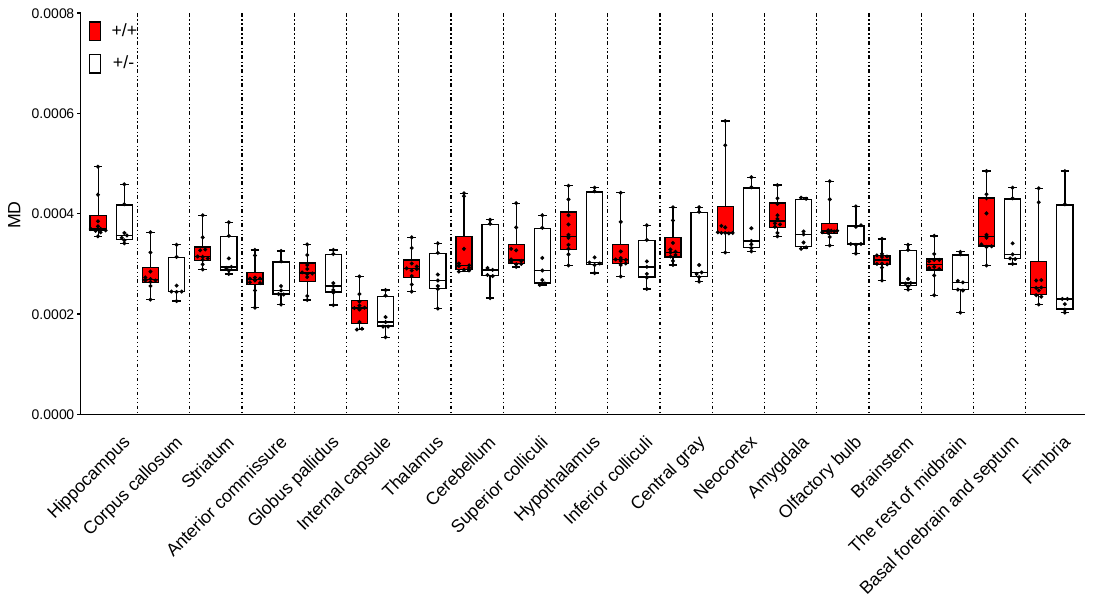

**B**
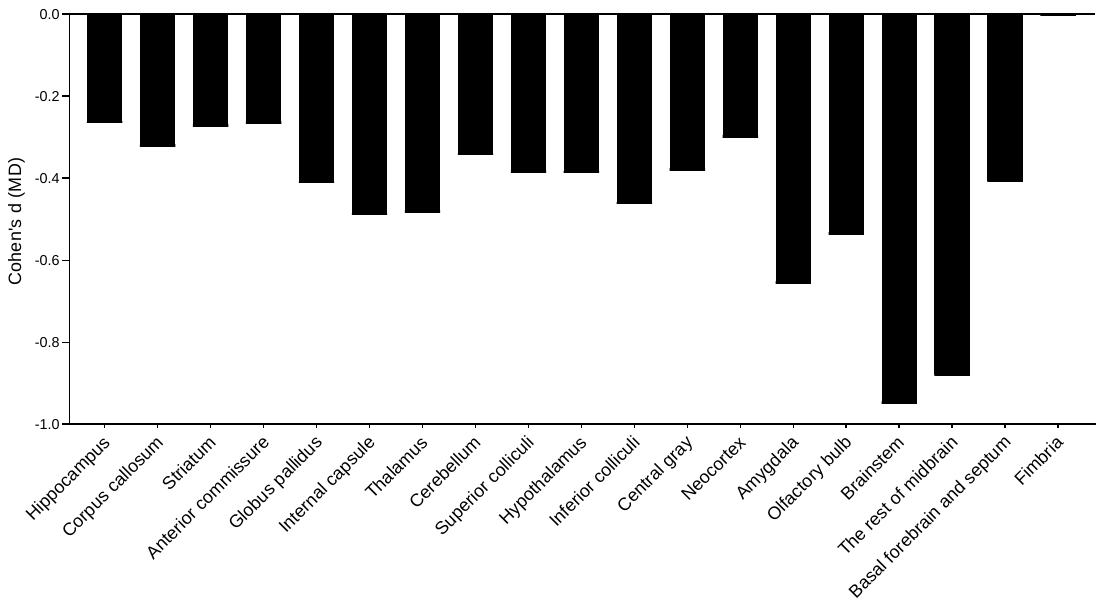

**Figure S5.** **(A)**. Mann-Whitney U-tests revealed no between-genotype differences in mean diffusivity (MD) values in any of the regions examined (all p > 0.05). **(B)**. Effect sizes of the MD differences between +/+ and +/- mice, as determined by Cohen’s d values.

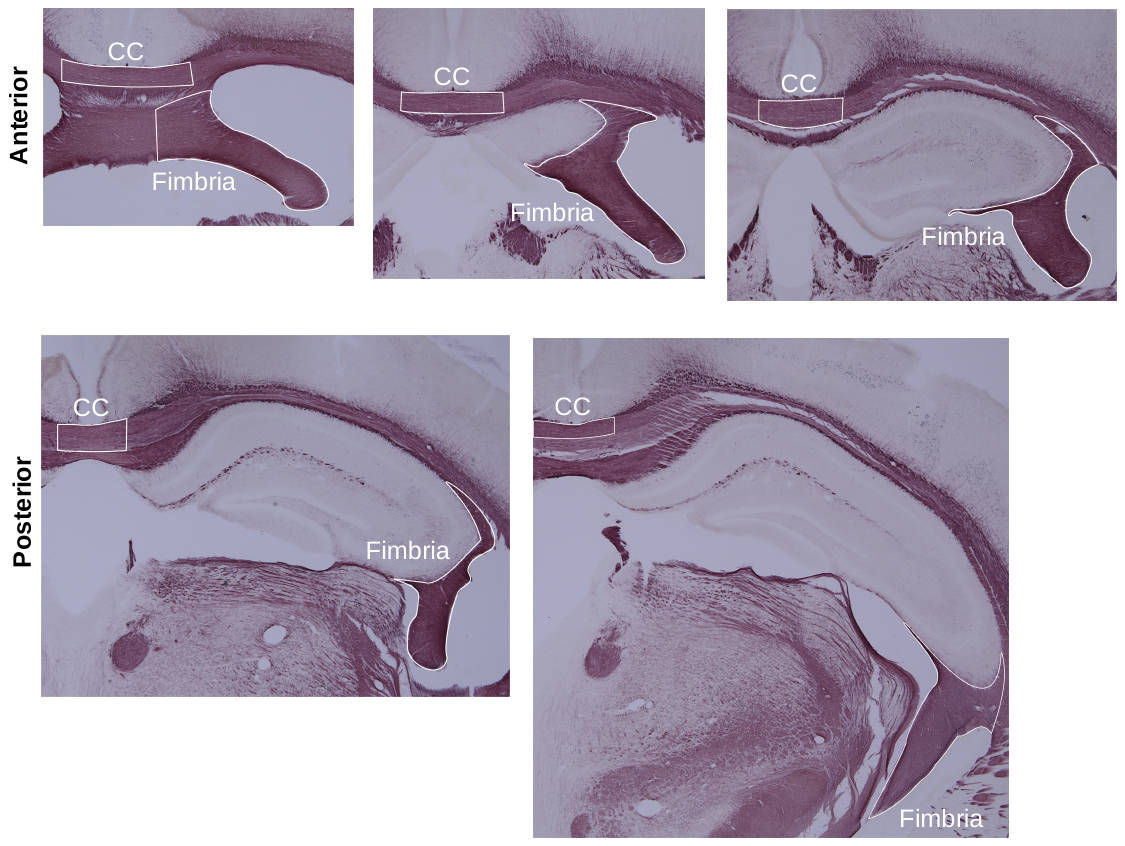
 **Figure S6.** Representative images of the anterior (Bregma –0.82 to –1.22mm) and posterior (Bregma –1.34 to –1.94mm) regions of the fimbria and corpus callosum (CC). This anterior-posterior division was based on areas exhibiting significant reductions in fractional anisotropy (FA) signals without necessarily following the anatomical definition of the antero-posterior division. Gold staining intensity was measured in the demarcated areas.

**A**

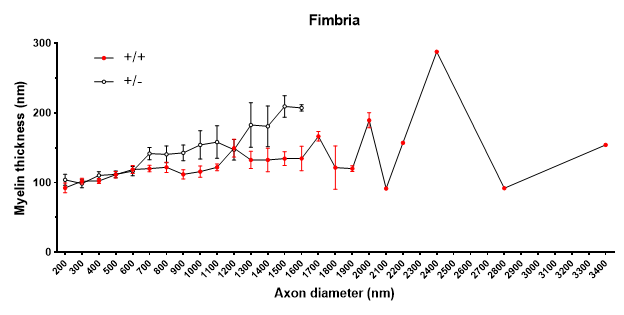

**B**
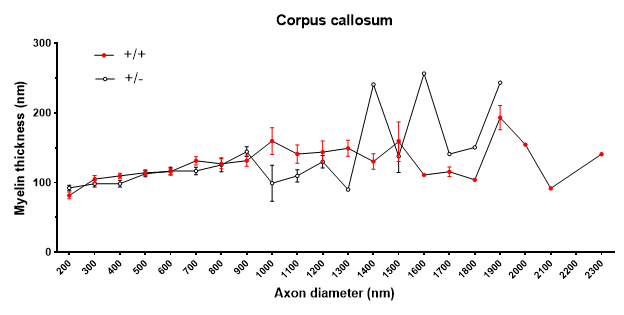
**Figure S7.** Average (± standard error of the mean [SEM]) myelin thickness at each 100-nm axon diameter unit. **(A)** In +/- mice, we observed increased myelin thickness for axons in the fimbria with diameters between ≥700 nm and <1,700 nm, and there were no axons of ≥1,700 nm in this region. **(B)** The same mice exhibited decreased myelin thickness for axons in the corpus callosum with diameters between ≥1,000 nm and <1,400 nm diameters, as well as increased myelin thickness for corpus callosum axons with diameters at and above 1,400 nm. Axons with diameters of ≥2,000 nm were not observed in the corpus callosum of +/- mice.

**A**
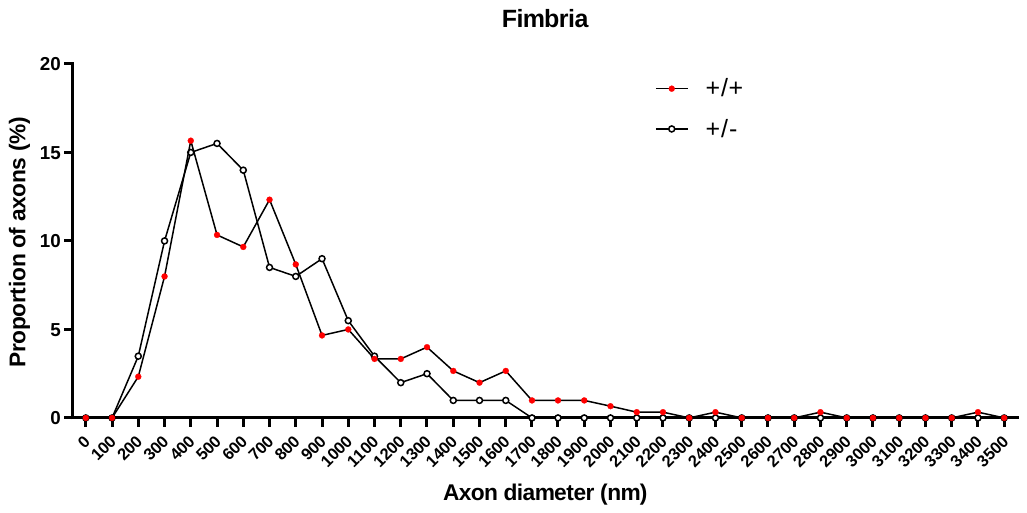

**B**
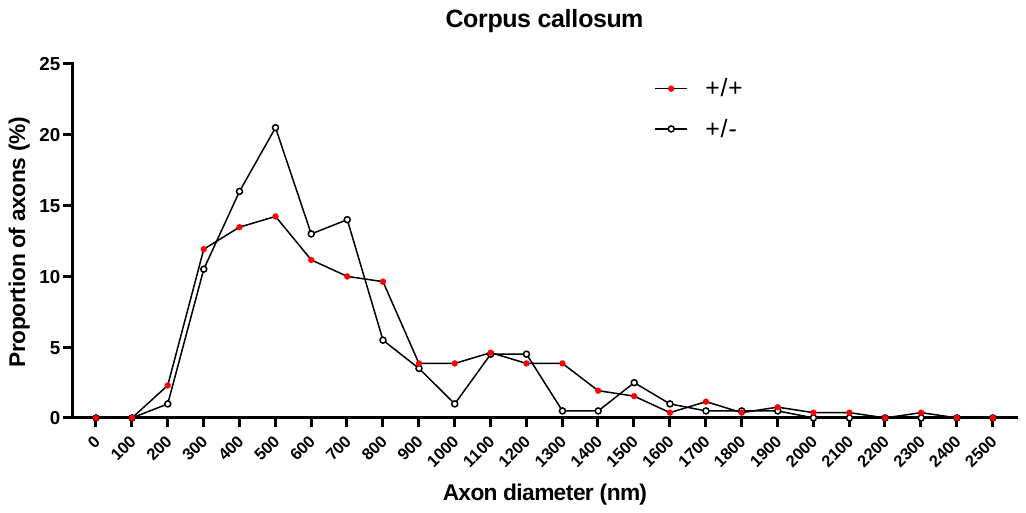
**Figure S8 .** The proportion of myelinated axons at each 100-nm axon diameter unit. **(A)** The relative proportion of myelinated axons in the fimbria with diameters between ≥500 and <1,200 nm was higher in +/- mice than in +/+ mice, while that of axons with diameters pf ≥1,200 nm was lower. No axons with diameters equal to or larger than 1,700 nm were observed in the fimbria (see also **Table S1**, Fimbria). **(B)** In the corpus callosum, the relative proportion of myelinated axons with diameters between ≥400 and <800 nm was higher in +/- mice than in +/+ mice, while that of axons with diameters equal to or larger than ≥800 nm was lower (see also **Table S1**, Corpus callosum). Values are expressed as percentages [(# of axons in each unit/# of all axons in region) x 100].

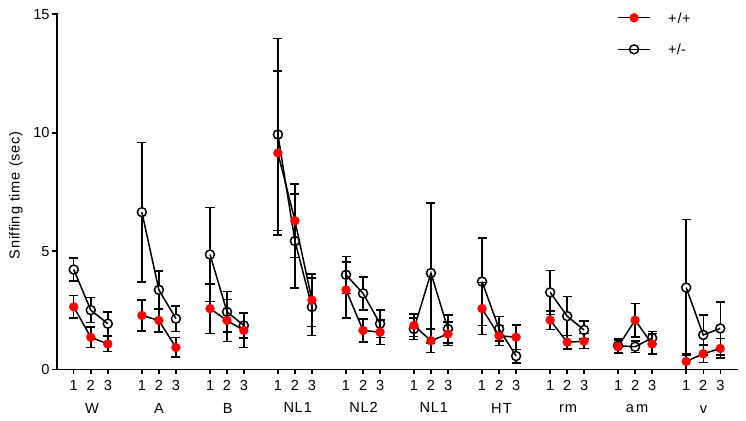

**Figure S9.** Olfactory responses to non-social and social odorants. The mean (± standard error of the mean [SEM]) sniffing time (s) at an Eppendorf tube containing each odorant. The assumptions of normality and homogeneity of variance were violated, as assessed using Shapiro–Wilk tests and Levene’s tests, respectively. Non-parametric Mann–Whitney U-tests, adjusted by Benjamini–Hochberg’s correction, revealed no differences between +/+ and +/- mice for any odorant in any session. W, water; A, almond odor; B, banana odor; NL, urine of non-littermate C57BL/6J male mouse; +/-, urine of a non-littermate +/- female mouse; rm, urine of the dam of tested mice, am, urine of a non-dam mother; v; urine of a virgin female C57BL/6 mouse. +/+, N = 14 and +/-, N = 14 for water; almond and banana, NL1, NL2, and HT. +/+, N = 26, +/-, N = 24 for rm and am, +/+, N = 9, +/-, N = 11 for v.

**Table S1**. Number (percentage) of axons in each axon diameter range (nm)

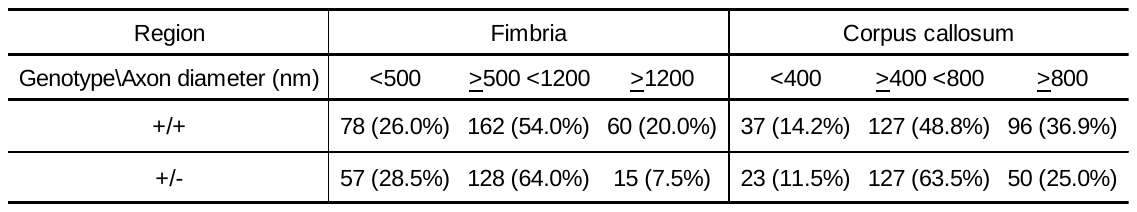

There were proportionally fewer myelinated axons of ≥1,200 nm in the fimbria (χ^2^(2, N=500) = 14.847, p=0.001) and fewer myelinated axons of ≥800 nm corpus callosum (χ^2^(2, N=460) = 10.106, p=0.006) of +/- mice than in those of +/+ mice. However, +/- mice exhibited more myelinated axons with diameters between ≥500 nm and <1,200 nm in the fimbria and between ≥400 nm and <800 nm in the corpus callosum than +/+ mice. Axon diameters were divided into three groups based on quantitative differences between +/+ and +/- mice (see **Fig. S8**). Fimbria, +/+, N=300, +/-, N=200; Corpus callosum, +/+, N=260, +/-, N=200.

|  | **Table S2. Intensity calculations** |  |
| --- | --- | --- |
| **Step** | **Value** | **Formula** |
| 1 | Brightness of target | tB/A |
| 2 | Brightness of negative control | ncB/A |
| 3 | Ratio of each negative control | (max ncB/A)/(ncB/A)=R |
| 4 | Adjusted brightness | tN/A^adj^=(tB/A)*R |
| 5 | Conversion to staining intensity | 1/(tB/A^adj^) |

B, Brightness of the target (t) region; A, Area of the target region; tB/A, brightness

per target area; ncB/A, Brightness per area unit of the cortex as a negative control (nc)

of non-specific staining; R, the corrective ratio for adjustment based on the intensity of

non-specific staining. The brightest cortical section (max ncB/A) was considered to be devoid of non-specific staining. Cortical areas exhibiting less brightness (<max ncB/A) were considered to exhibit a degree of non-specific staining, resulting in R values larger than 1.0. If tB/A is multiplied by R, the adjusted brightness value ((tB/A)*R ) is corrected proportional to the degree of non-specific staining. As this value is negatively proportional to the degree of gold staining, we used its inverse value to represent the degree of gold staining.

| **Table S3. Assay ID numbers** | |
| --- | --- |
| **Gene Symbol** | **Assay ID** |
| *Tbx1 exon 2-3* | Mm01342798_m1 |
| *Ng2 (Cspg4)* | Mm00507257_m1 |
| *Pdgfr2* | Mm00440701_m1 |
| *MBP* | Mm01266402_m1 |
| *MOG* | Mm01279062_m1 |
| *Cyc1* | Mm00470540_m1 |

**Table S4. Dimensions and exemplar combinations used for attentional set shifting**

| Task | Dimension | | Exemplar combinations | |
| --- | --- | --- | --- | --- |
|  | Relevant | Irrelevant | Correct | Incorrect |
| SD  CD    IDS I    IDS II    IDS III    IDS IV    IDS IV rev  EDS | Odor (O) Medium (M)  Odor    Odor  Odor    Odor    Odor  Odor  Medium | ---------    Medium  Medium  Medium  Medium  Medium  Medium  Odor | O1&M1  O1&M1  O1&M2  O3&M3  O3&M4  O5&M5  O5&M6  O7&M7  O7&M8  O9&M9  O9&M10  O10&M9  O10&M10  M11&O11  M11&O12 | O2&M2  O2&M2  O2&M1  O4&M4  O4&M3  O6&M6  O6&M5  O8&M8  O8&M7  O10&M10  O10&M9  O9&M10  O9&M9  M12&O12  M12&O11 |

See **S-Table 5** for each exemplar. SD, simple discrimination; CD, compound discrimination;

IDS, intra-dimensional shift; rev, reversal; EDS, extradimensional shift.

**Table S5: Odorants and media used as exemplars of attentional set shifting**

| **Pair** | **Test** | **Exemplar** | **Odor** | **Medium** |
| --- | --- | --- | --- | --- |
| 1 | SD, CD | 1 | Sage（O1） | Alpha drip（M1） |
|  |  | 2 | Cinnamon （O2） | Paper chip（M2） |
| 2 | IDS I | 3 | Coriander （O3） | Carefresh Natural（M3） |
|  |  | 4 | Onion （O4） | Kaykob bedding（M4） |
| 3 | IDS II | 5 | Garlic （O5） | Eco bedding（M5） |
|  |  | 6 | Paprika （O6） | Sphang moss（M6） |
| 4 | IDS III | 7 | Rosemary （O7） | Aspen Bedding（M7） |
|  |  | 8 | Cloves （O8） | Aquarium Gravel（M8） |
| 5 | IDS IV, IDSIV rev | 9 | Thyme （O9） | Rapti bark（M9） |
|  |  | 10 | Black Pepper（O10） | Shredded paper（M10） |
| 6 | EDS | 11 | Cumin （O11） | ExquisiCat（M11） |
|  |  | 12 | Cardamom（O12） | Carefresh Ultra（M12） |

SD, simple discrimination; CD, compound discrimination; IDS, intra-dimensional shift;

rev, reversal; EDS, extradimensional shift.
